# A Scalable Adaptive Quadratic Kernel Method for Interpretable Epistasis Analysis in Complex Traits

**DOI:** 10.1101/2024.03.09.584250

**Authors:** Boyang Fu, Prateek Anand, Aakarsh Anand, Joel Mefford, Sriram Sankararaman

**Affiliations:** Department of Computer Science, UCLA, Los Angeles, CA, USA; Semel Institute for Neuroscience and Human Behavior, UCLA, Los Angeles, CA, USA; Department of Human Genetics, David Geffen School of Medicine, UCLA, Los Angeles, CA, USA; Department of Computational Medicine, David Geffen School of Medicine, UCLA, Los Angeles, CA, USA

## Abstract

Our knowledge of the contribution of genetic interactions (*epistasis*) to variation in human complex traits remains limited, partly due to the lack of efficient, powerful, and interpretable algorithms to detect interactions. Recently proposed approaches for set-based association tests show promise in improving power to detect epistasis by examining the aggregated effects of multiple variants. Nevertheless, these methods either do not scale to large numbers of individuals available in Biobank datasets or do not provide interpretable results. We, therefore, propose QuadKAST, a scalable algorithm focused on testing pairwise interaction effects (also termed as *quadratic effects*) of a set of genetic variants on a trait and quantifying the proportion of phenotypic variance explained by these effects.

We performed comprehensive simulations and demonstrated that QuadKAST is well-calibrated. Additionally, QuadKAST is highly sensitive in detecting loci with epistatic signal and accurate in its estimation of quadratic effects. We applied QuadKAST to 53 quantitative phenotypes measured in ≈ 300, 000 unrelated white British individuals in the UK Biobank to test for quadratic effects within each of 9, 515 protein-coding genes (after accounting for linear additive effects). We detected 32 trait-gene pairs across 17 traits that demonstrate statistically significant signals of quadratic effects (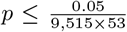 accounting for the number of genes and traits tested). Our method enables the detailed investigation of epistasis on a large scale, offering new insights into its role and importance.

## 1 Introduction

Genome-wide association studies (GWAS) have revolutionized the field of human genetics by providing valuable insights into the genetic basis of complex traits and diseases. The primary goal of GWAS is to identify statistically significant associations between specific genetic variants and the phenotype being studied. During the past decades, models based on additive assumptions have been successfully applied to identify variants that impact complex traits and diseases [1–4] due to their simplicity and interpretability. Recent studies indicate that interaction effects between genes or genetic variants that go beyond mere additivity can play an overlooked role in shaping complex traits [5]. Such interactions have been proposed as key factors in both human complex trait variation and disease susceptibility [6]. Epistasis also potentially accounts for some of the “missing heritability” not explained by additive genetic factors alone [7, 8]. Despite its likely importance, efficient methods for detecting, dissecting, and interpreting complex epistatic interactions remain largely undeveloped, and our knowledge of the epistasis remains limited [9]. Having an efficient way of identifying and understanding epistasis could greatly advance our understanding of underlying biological pathways [10, 11] and can potentially increase the generalizability of polygenic scores within [12] and across different ancestral populations [13, 14].

Despite its importance, characterizing the role of epistasis in complex traits presents several challenges. The task of examining all potential interactive relationships among SNPs and genes necessitates navigating a large feature space that expands exponentially with the increasing order of interactions. A number of methods have been developed to search [15–17] for pairs of genetic variants that show evidence for epistatic effects from a large combinatorial space. However, such approaches have low statistical power due to the stringent thresholds needed to account for the number of tests performed. As a result, successful detection of epistatis requires examining a large number of individuals to obtain adequate power.

An alternative approach to identify trait-relevant genetic variants focuses on grouping variants into “sets” and jointly estimating the effects of all variants within each set [1, 18–20]. By reducing the number of statistical tests performed and hence the multiple testing burden, these methods can obtain increased power over approaches that aim to identify individual variants. Moreover, by aggregating relevant signals into sets, this strategy can enhance the statistical power for detecting the relevant signals. Existing set-based tests have shown their efficacy for detecting associations between complex traits and sets of rare and common variants [21–23]. However, these approaches, while largely scalable, focus primarily on testing the additive effect of variants within a set. None of the existing approaches [1, 20] can test epistatic effects in largescale biobanks. One approach to test for non-linear effects relies on the “kernel trick” that enables implicit computation of the inner product of potentially high-dimensional non-linear transformations of the input genotypes. However, this workaround does not alleviate the computational burden with the key bottleneck being the eigendecomposition of the *N × N* kernel matrix which has a 𝒪 (*N* ^3^) complexity, making the computation infeasible on large-scale data. Recent works, such as FastKAST, have ameliorated the computational challenge by employing advanced sampling strategies [24] to approximate the kernel decomposition. These methods enable the tests for epistasis within sets in an efficient manner, with a time complexity of 𝒪 (*ND*^2^) [5], and exhibit enhanced testing power when extending the hypotheses beyond additive models. However, these approaches can only approximate shift-invariant kernels, such as the radial basis function kernels, which excludes quadratic and general polynomial kernels, and hence lack the interpretability associated with testing for pairwise interactions among SNPs. The lack of flexibility in the kernel design makes it difficult to interpret the results. Overall, we lack efficient yet interpretable methods to identify and quantify the epistasis effects within sets of genetic variants.

We propose a novel algorithm, **Quad**ratic **K**ernel-based **AS**sociation **T**est (QuadKAST), to address the major limitations of existing set-based association test approaches. Unlike existing approaches, QuadKAST aims to test for the effect of pairwise genetic interactions. This approach offers several advantages. First, the pairwise effects offer an interpretable (and the simplest) model of epistasis. Second, besides merely performing epistasis testing (a binary answer for statistical significance), QuadKAST estimates the proportion of phenotypic variance explained by non-linear effects.

We performed comprehensive simulations and demonstrated the good calibration and power of QuadKAST. Compared to existing methods, QuadKAST is the only method that is feasible to run on Biobank-scale datasets with hundreds of thousands of individuals. QuadKAST offers scalable and calibrated set-based tests of pairwise epistatic effects and estimates the variance components associated with these effects. We applied QuadKAST to test for epsitatic effects in protein-coding genes for 53 quatitative traits measured in the UK-Biobank to identify 32 trait-genes pairs demonstrating significant epistatic signals (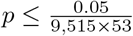 accounting for the number of genes and traits tested). Our method enables systematic investigation of epistasis on a large scale, offering new insights into its role and importance.

## 2 Methods

### Set-based association testing of linear additive genetic effects

Consider a *N × M* matrix representing the standardized genotypes of *N* individuals at *M* SNPs: 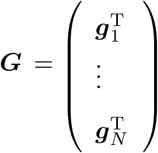where ***g***_*n*_, *n* ∈ *{*1, …, *N}* is the vector of genotypes at *M* SNPs for a specific individual *n*. Let ***y*** ∈ ℝ^*N*^ represent the phenotypes across *N* individuals and ***X*** represents an *N × K* matrix of covariates.

In the context of set-based association testing, the matrix ***G*** is typically constructed by aggregating variants within a genomic region while matrix ***X*** incorporates factors such as sex, age, and genetic principal components (PCs) to account for population structure. The objective of set-based association testing is to ascertain whether the variants within the defined set exhibit, in aggregate, association with the trait ***y*** where the association is typically assumed to be linear and additive. Formally, we assume that each of the *M* SNPs within the set independently and additively contributes to the trait with effect sizes drawn from a normal distribution resulting in the following model.

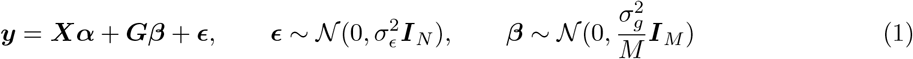

Here 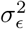 denotes the residual or noise variance while 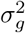 represents the proportion of phenotypic variance explained by additive genetic effects at the SNPs considered. ***α*** denotes the fixed effects associated with the covariates. The objective of set-based association testing is formulated as a test of the hypothesis 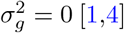.

### Quadratic association test

Beyond a linear additive relationship, the genetic variants within a set may modulate the trait through their interactions with each other. We can expand upon the model in Equation 1 to include nonlinear associations between ***G*** and ***y*** through the use of a feature map *ϕ* : ℝ^*M*^ → ℝ^*D*^. This map transforms the vector of genotypes at *M* SNPs into a *D*-dimensional vector. While the feature map *ϕ* could represent an arbitrary non-linear function, considerations of interpretability lead us to restrict *ϕ* to functions that capture pairwise interactions across the *M* SNPs (quadratic feature maps). We can define two such quadratic feature maps depending on whether we allow for self-interactions at a SNP or not:

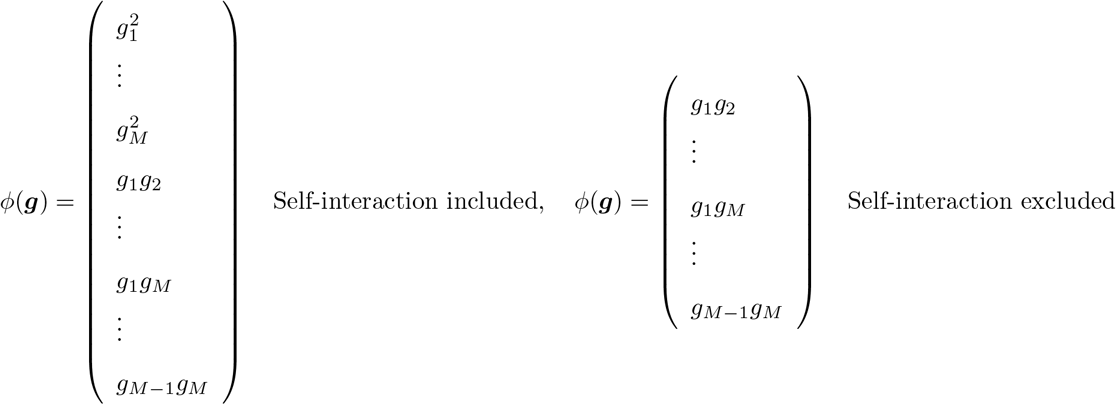

We now model the aggregate effect of pairwise genetic interactions on the phenotype, termed *quadratic effects*, as:

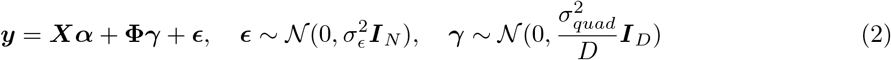

Here 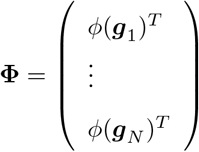 while ***γ*** ∈ ℝ^*D*^ is a random vector of effects associated with each pairwise interaction. 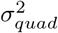 represents the variance attributed to all pairwise effects or quadratic effects across the set of SNPs (*quadratic variance component*). To ensure that this model is sensitive to non-additive genetic effects, we include additive genetic effects (represented by the matrix ***G***) within ***X*** (effectively regressing out their contribution to the phenotype).

Integrating out the random effects ***γ***, the distribution of ***y*** follows 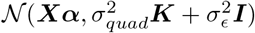. Here ***K*** is a *N × N* kernel matrix where *K*_*i,j*_= *ϕ*(***g***_***i***_)^*T*^ *ϕ*(***g***_***j***_)*/D* so that 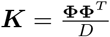.

In this model, we aim to answer two primary questions. First, we want to test whether the phenotypic value is associated with the aggregate pairwise interactions effects across the SNPs within the set of interest (described by Φ), *i*.*e*., we aim to test the hypothesis 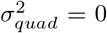. Second, if 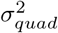 is nonzero, we would like to estimate the variance component parameter 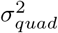. Importantly, we would like to develop procedures for hypothesis testing and variance component estimation that can be applied to large-scale biobanks where the number of individuals *N* is large (of the order of hundreds of thousands).

### Hypothesis test

The hypothesis of interest is whether the variance explained by the pairwise interaction effects of the target set of variants 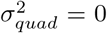, conditioning on the additive effect of the variants and other covariates. Previous work has shown that including the genetic variants in the window surrounding the target set as fixed-effect covariates ensures that the additive effects are residualized from the phenotype [5]. We, therefore, adopt this strategy in our work. To test the hypothesis that 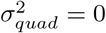, we define the score test statistic:

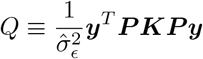

where 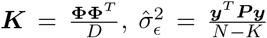, and ***P*** = (***I*** − ***X***(***X***^*T*^ ***X***)^−1^***X***^*T*^) is the projection matrix with ***X*** as the covariates matrix. See Supplementary Note S1.1 for derivation of the score test statistic. Under the null hypothesis, previous work [5] has shown that the distribution of the score test statistic is:

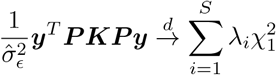

where *λ*_*i*_ is the *i*^*th*^ eigenvalue of ***PKP*** and *S* is the rank of ***PKP***. However, the construction of ***K*** and the eigen-decomposition of ***PKP*** scales as *𝒪* (*N* ^2^*D*) and *𝒪* (*N* ^3^) which does not scale with to datasets with a large number of individuals (*N*).

To overcome this bottleneck, we first compute the singular value decomposition of ***P* Φ** so that the eigenvalue *λ*_*i*_ can be computed from the corresponding singular values. The singular values of ***P* Φ** can be computed in 𝒪 (*ND*^2^) time (for *D < N*) leading to an efficient algorithm for large numbers of individuals provided the number of SNPs in the set is not too large (since *D* = 𝒪 (*M* ^2^)).

### Variance component estimation

#### No covariates

Let us first consider the setting where there are no covariates included in the model. The distribution of the phenotype can be written as:

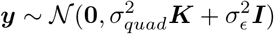

The variance components 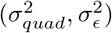 can be estimated by maximizing the log-likelihood:

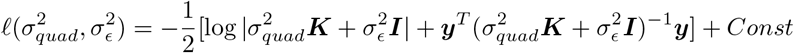

It can be shown that the likelihood can be maximized by maximizing the profile log-likelihood function obtained by maximizing or profiling out 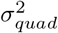 from the log-likelihood and writing it in terms of 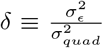 (Supplementary Note S1.2):

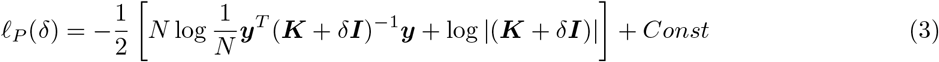

However, naively evaluating and optimizing the profile log-likelihood function involves an iterative algorithm with 𝒪 (*N* ^3^) time complexity in each iteration rendering this approach impractical when the number of individuals increases.

Given the eigen-decomposition of ***K*** = ***U*** *diag*(*ρ*_1_, …, *ρ*_*N*_)***U*** ^*T*^ where ***U*** ∈ ℝ^*N×N*^ is the matrix of eigenvectors and (*ρ*_1_, …, *ρ*_*N*_) are the eigenvalues of ***K***, Equation 3 can be rewritten as

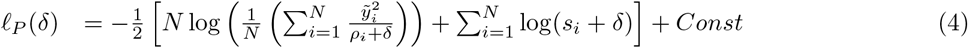

Here 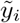 is the *i*^*th*^ entry of 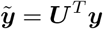. With this transformed representation, the profile log-likelihood function can be optimized with 𝒪 (*N*) time complexity in each iteration once the eigen-decomposition of ***K*** and the transformed phenotype 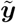 have been computed.

In our application, the matrix 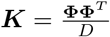 for **Φ** ∈ ℝ^*N×D*^ where *D* = 𝒪 (*M* ^2^) *< N*. As a result, the rank (*R*) of ***K*** is lower than its dimensionality. This allows us to further rewrite the profile log-likelihood as:

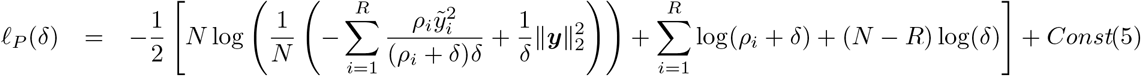

Evaluating *ℓ*_*P*_ in this setting requires computing 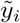 and *ρ*_*i*_, *i* ≤ *R* which can be obtained by a one-time computation of the *R* non-zero eigenvalues and corresponding eigenvectors of ***K***. Computation of these eigenvalues and eigenvectors can be obtained in 𝒪 (*ND*^2^) time from a singular value decomposition (SVD) of **Φ**) while subsequent evaluation of *ℓ*_*P*_ requires 𝒪 (*D*) time (see Supplementary Note S1.2). We used the built-in Scipy [25] package “Nelder-Mead” algorithm [26] for the optimization.

#### Including covariates

When covariates are incorporated into the model, the phenotypic distribution can be expressed as

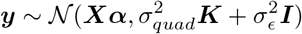

In this setting, the profile restricted log-likelihood function can be computed efficiently when ***K*** is low-rank (Supplementary Note S1.3):

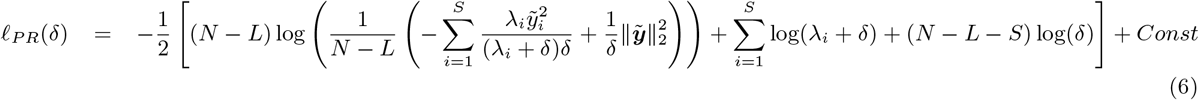

Here 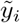 is the *i*^*th*^ entry of the transformed phenotype: 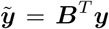 where ***B*** is the matrix of eigenvectors of ***PKP*** with *S* non-zero eigenvalues *λ*_1_, *λ*_2_, …, *λ*_*S*_. To compute 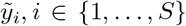, we need to compute eigenvectors (columns of ***B***) corresponding to the non-zero eigenvalues of ***PKP*** which is equivalent to the corresponding left singular vectors of ***P* Φ**. Finally, *ρ*_*i*_, *i* ∈ *{*1, …, *S}* are the non-zero eigenvalues of ***PKP*** which are also obtained from the corresponding singular values of ***P* Φ**. Both these quantities can be obtained in 𝒪 (*NKD* + *NK*^2^ + *K*^3^ + *ND*^2^) time, where *K* is the columns number of ***X***. Thus, the profile restricted log-likelihood as represented in Equation 18 can be optimized with a 𝒪 (*NKD* + *NK*^2^ + *K*^3^ + *ND*^2^) one-time computation followed by *O*(*D*) time to evaluate this function subsequently.

#### Standard error estimation

Once the parameters, 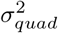 and 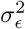, have been estimated, we can further compute the corresponding standard error using the observed Fisher information matrix evaluated at the MLE or the maximum restricted likelihood (REML) estimates.

## 3 Results

### Datasets

We obtained a set of *N* = 291, 273 unrelated white British individuals measured at *M* = 459, 792 common SNPs genotyped on the UK Biobank (UKBB) Axiom array to use in simulations by extracting individuals that are *>* 3rd-degree relatives and excluding individuals with putative sex chromosome aneuploidy. Unless otherwise specified, all simulations and real data analysis were conducted using this dataset.

By default, the sets were defined by the start and end physical position of protein-coding genes with the number of SNPs ≥ 3 and ≤ 50 resulting in 9, 515 sets of SNPs from protein-coding genes. In real data tests, we included sex, age, and the top 20 genetic principal components (PCs) as covariates in our analysis.

### Covariates and phenotypes

We selected 53 quantitative traits that were measured in the UKBB and which had been analyzed in prior studies of non-linear genetic effects [27, 28] (see Supplementary table S1). All traits were transformed using inverse rank-normalization. We included sex, age, and the top 20 genetic principal components (PCs) as covariates in our analysis for all phenotypes. We used PCs computed in the UKBB from a super-set of 488, 295 individuals. Extra covariates were added for diastolic/systolic blood pressure (adjusted for cholesterol-lowering medication, blood pressure medication, insulin, hormone replacement therapy, and oral contraceptives) and waist-to-hip ratio (adjusted for BMI).

### Calibration of QuadKAST

First, we assessed QuadKAST in terms of controlling type-I error by applying it to simulated data in the absence of interactive effects. We simulated phenotypes based on genotypes of SNPs on the UKBB array across unrelated white British individuals in the UK Biobank (*N* = 291, 273 individuals, *M* = 459, 792 SNPs). For computational reasons, we randomly sub-sampled 50K individuals. We performed simulations under four genetic architectures with additive but no interaction effects: infinitesimal model (causal variants ratio = 1); non-infinitesimal model (causal variants ratio = 0.001), which is further divided into three categories, each having a different MAF range for the causal variants from the three possible ranges: [0.01, 0.05] (RARE), [0.05, 0.5] (COMMON), [0, 0.5] (ALL). In all settings, the additive heritability was fixed at 0.5. We applied QuadKAST on sets of SNPs typed on the UKBB array where each set is one of 9, 515 protein-coding genes (genes with *>* 50 SNPs excluded). We observed that QuadKAST is well-calibrated across all the simulation settings (Figure 1a).

**Figure 1:**
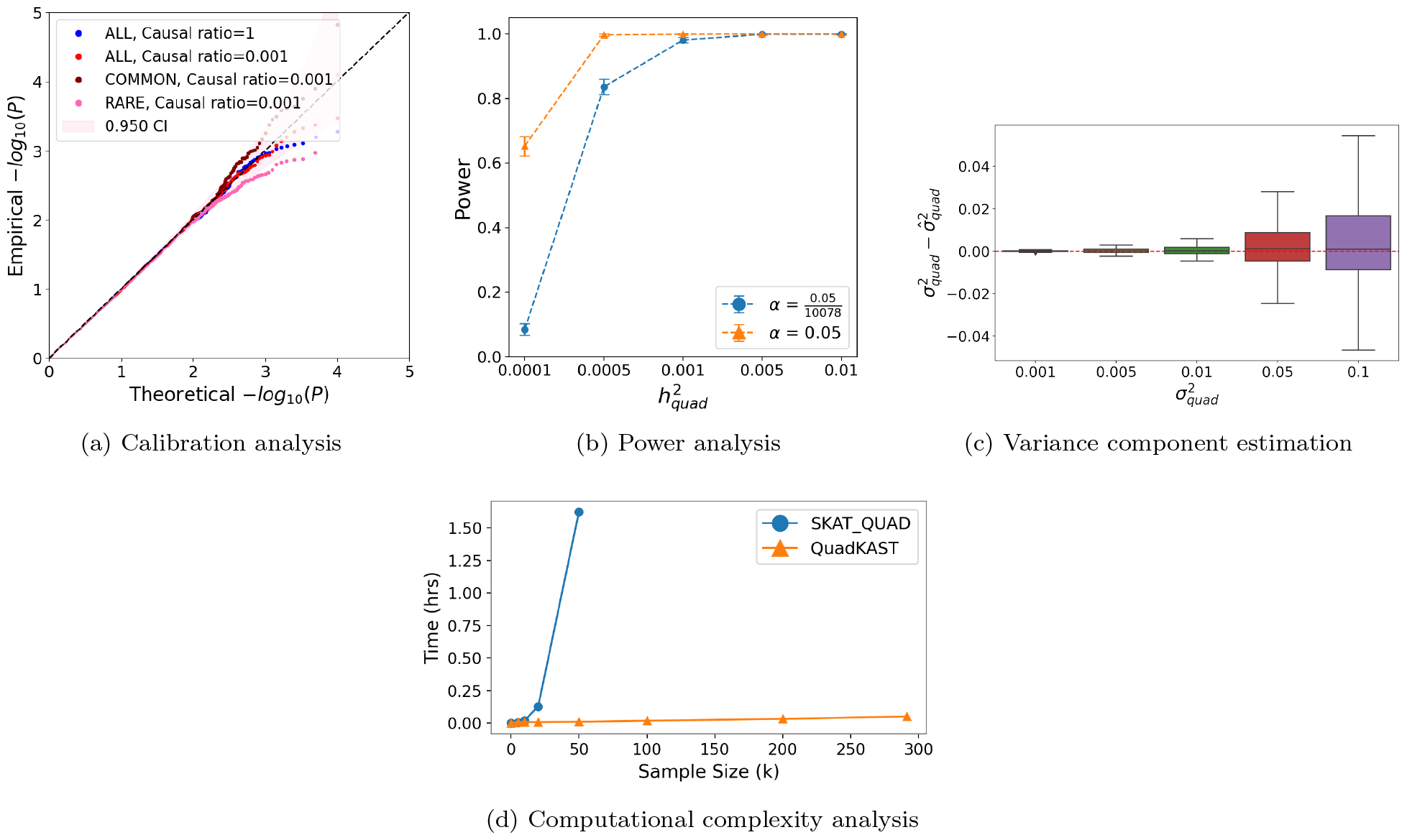
(a) **Calibration**. We applied QuadKAST to SNPs within 9, 515 protein-coding genes for four genetic architectures that consist entirely of linear additive effects (N = 50K individuals, UK Biobank (UKBB) array data) (b) **Power analysis**. We simulated traits with varying quadratic variance component on a randomly selected subset of 5K individuals from unrelated white British individuals in UKBB. We applied QuadKAST to 1, 000 randomly selected protein-coding genes gene and defined power as the ratio of p-values reported by QuadKAST that pass the significance threshold *α*. (c) **Accuracy**. Similar to (b), we applied QuadKAST to estimate the quadratic variance component at each gene. (d) **Runtime**. We compared the runtime of SKAT and QuadKAST by fixing the number of SNPs *M* = 100 and varying the number of individuals (average runtime across 10 replicates).

One of the features of QuadKAST is a flexible choice of the quadratic kernel that is used to test for interactions. The default quadratic kernel function encodes all pair-wise interactions within a set (with variants that depend on whether self-interactions are included). We confirmed calibration of quadratic kernel functions with and without the inclusion of self-interactions, termed *self-interaction included* and *self-interaction excluded* respectively (Supplementary Figure S2). We further confirmed calibration for lower p-value thresholds (*α* = 1 *×* 10^−6^) using a large number of tests on one exemplar genetic architecture (ALL, Causal ratio=0.001) (Supplementary Tables S2, S3).

### Power Analysis of QuadKAST

Our next set of experiments sought to evaluate the power of QuadKAST. Specifically, our generative model is

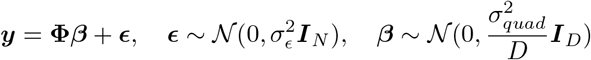

In these settings, the self-interactions contribute to the phenotypic variance. We varied 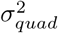 and for each parameter setting, we randomly selected 1, 000 sets from the set of 9, 515 protein coding genes and a random set of 5K unrelated, white British individuals. Then we applied QuadKAST to test 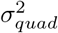 in these sets with power defined by the estimated p-value passing the significance threshold *α*. QuadKAST can achieve power over 0.8 with the Bonferroni-corrected significance threshold at a signal strength 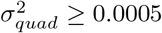 (Figure 1b). A subsequent power comparison, contrasting the self-interaction excluded setting during both the simulation process and the variance component estimation process showed that QuadKAST is powerful in detecting the signal in both scenarios (Supplementary Figure S4).

### Accuracy of QuadKAST Variance Component Estimation

Beyond statistical power, we aimed to test the accuracy of the variance component estimates from QuadKAST using the same simulation setup as we did in power analysis. QuadKAST can accurately estimate the variance component (Figure 1c with a more comprehensive analysis of accuracy with respect to sample size, feature size, and signal strength in Supplementary Figure S3). We also find that the variance components are accurately estimated with and without the inclusion of self-interactions (Supplementary Figure S5).

### Computational Efficiency of QuadKAST

Finally, we compared the runtime of QuadKAST and the quadratic kernel option in the popular SKAT software (SKAT QUAD) [29]. We chose a set consisting of 100 SNPs corresponding to the 99^*th*^ percentile of the distribution of UKBB array SNPs contained within protein-coding genes (Supplementary Figure S1). We then varied the sample size and profiled the runtime and memory usage of QuadKAST and SKAT QUAD (with a pre-defined limit of 4 hours for the runtime). While the average runtime of QuadKAST on a single set is under 5 minutes for UK Biobank size data (≈ 300k), SKAT QUAD reaches the time limit with larger than 100k samples (Figure 1d). Running SKAT QUAD on UK Biobank size data would take more than 10 hours, not to mention the memory consumption of constructing the *N × N* kernel matrix (Supplementary Figure S6b). Based on these experiments, it is evident that running SKAT QUAD on UK Biobank scale data would be infeasible due to computational and memory limitations while QuadKAST is about 100x faster.

### Application of QuadKAST to UK Biobank phenotypes

After confirming the calibration and power of QuadKAST, we applied it to 53 quantitative traits in the UK Biobank measured across *N* = 291, 273 unrelated white British individuals and to SNPs on the UKBB array grouped into sets defined by 9, 515 genes (see Section 3 for more details on the data preparation). Before testing the pairwise interaction across variants in each gene on the quantitative traits, we first regressed out the additive effect of the SNPs around the target set (the surrounding sets two times larger than the target set in addition to the target set), together with covariates that include 20 PCs, age, and sex information on the target traits. To account for SNPs that are in high LD, we transformed the matrix of additive genotypes using a SVD before running linear regression. We included the self-interaction when testing the quadratic effect of each set (unless mentioned otherwise).

With this strategy, we identified 32 trait-gene pairs to be statistically significant 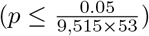 across 17 quantitative traits after accounting for the total genes and traits tested (Table 1). We estimated the quadratic variance component across all 32 significant trait-gene pairs to observe that the median ratio 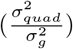 is 0.15. In addition, the influence of the epistatic effect may be significantly greater than the additive effect for specific pairs of genes and traits. For instance, the variance component ratio attributed to the quadratic effect is more than 20 times higher than that of the additive effect for the PRG3 gene on the Eosinophil count traits. We also note that the epistatic effect of LPA for trait Lipoprotein-A is consistently large 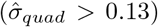. Our results highlight the significance of epistasis in complex trait variation (Supplementary table S4).

**Table 1:** Significant epistatic trait-gene pairs. Trait-gene pairs with statistically significant epistatic effects (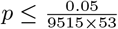 accounting for the number of genes and traits tested) were reported. The −*log* p-values were reported in entry −*log*_10_ Pval, with a precision level bounded by 13, using default settings. We further compared them with results where the top 40 PCs are considered as the covariates, denoted by −*log*_10_ *Pval (40 PC)* column, we report p-values when regressing 40 PCs. In the −*log*_10_ *Pval (Imputed)* column, we report p-values using UKBB imputed data.

| Trait | CHR | Gene | Start<br>(MB) | End<br>(MB) | $-\log_{10}$ Pval | $-\log_{10}$ Pval<br>(40 PC) | $-\log_{10}$ Pval<br>(Imputed) |
| --- | --- | --- | --- | --- | --- | --- | --- |
| Alanine aminotransferase | 22 | PNPLA3 | 44.32 | 44.34 | 15.18 | 15.18 | $\geq 13$ |
|  | 22 | SAMM50 | 44.35 | 44.39 | 8.02 | 7.97 | 6.61 |
| Apolipoprotein B | 19 | APOE | 45.41 | 45.41 | $\geq 13$ | $\geq 13$ | $\geq 13$ |
|  | 19 | BCAM | 45.31 | 45.32 | 9.03 | 9.02 | 2.84 |
| Aspartate aminotransferase | 22 | PNPLA3 | 44.32 | 44.34 | 11.92 | 11.91 | 14.63 |
| Creatinine | 1 | DNAJC16 | 15.86 | 15.89 | 7.05 | 7.07 | 6.42 |
| Direct bilirubin | 2 | SAG | 234.22 | 234.26 | 7.56 | 7.55 | 3.43 |
| | 2 | DGKD | 234.26 | 234.38 | $\geq 13$ | 14.81 | 7.02 |
| | 2 | USP40 | 234.39 | 234.47 | $\geq 13$ | $\geq 13$ | 4.51 |
| | 2 | UGT1A8 | 234.53 | 234.68 | $\geq 13$ | $\geq 13$ | $\geq 13$ |
| Eosinophil count | 11 | PRG3 | 57.14 | 57.15 | $\geq 13$ | $\geq 13$ | 5.17 |
| HDL cholesterol | 16 | CETP | 57.00 | 57.02 | 9.81 | 9.87 | 5.13 |
| Hemoglobin A1c | 10 | HK1 | 71.05 | 71.16 | 7.71 | 7.68 | 1.55 |
| LDL direct | 19 | APOE | 45.41 | 45.41 | 8.91 | 8.88 | 9.37 |
| Lipoprotein-A | 6 | IGF2R | 160.39 | 160.53 | $\geq 13$ | $\geq 13$ | 8.31 |
|  | 6 | SLC22A2 | 160.64 | 160.68 | 8.54 | 8.53 | 7.77 |
| | 6 | SLC22A3 | 160.77 | 160.87 | $\geq 13$ | $\geq 13$ | 4.13 |
| | 6 | LPA | 160.95 | 161.09 | $\geq 13$ | $\geq 13$ | $\geq 13$ |
|  | 6 | PLG | 161.12 | 161.17 | 8.54 | 8.54 | 12.11 |
| | 6 | AGPAT4 | 161.56 | 161.65 | $\geq 13$ | $\geq 13$ | 3.81 |
| Mean corpuscular hemoglobin | 22 | TMPRSS6 | 37.46 | 37.50 | 8.72 | 8.75 | 2.69 |
| Mean sphered cell volume | 1 | OR10Z1 | 158.58 | 158.58 | 7.39 | 7.40 | 7.71 |
|  | 1 | SPTA1 | 158.58 | 158.66 | 9.31 | 9.32 | 7.21 |
| Monocyte count | 22 | CECR1 | 17.66 | 17.69 | 8.80 | 8.80 | 2.97 |
| Platelet distribution width | 20 | TUBB1 | 57.59 | 57.60 | $\geq 13$ | $\geq 13$ | $\geq 13$ |
|  | 20 | EDN3 | 57.88 | 57.90 | 9.45 | 9.50 | 0.01 |
| SHBG | 17 | TNK1 | 7.29 | 7.29 | 7.37 | 7.35 | 2.42 |
| Urate | 1 | DNAJC16 | 15.86 | 15.89 | 8.26 | 8.27 | 11.42 |
| | 4 | SLC2A9 | 9.83 | 10.03 | $\geq 13$ | $\geq 13$ | $\geq 13$ |
| | 4 | WDR1 | 10.08 | 10.12 | 14.88 | $\geq 13$ | 14.42 |
|  | 4 | MEPE | 88.76 | 88.77 | 7.36 | 7.28 | 0.57 |
| Urea | 1 | MUC1 | 155.16 | 155.16 | 8.33 | 8.34 | 7.14 |

In the subsequent sections, we explored the robustness of our results. Specifically, we assessed the stability of our results to the population stratification and the potential missingness of the features using the imputed genotype dataset. Further, we investigate the contribution of individual interactions to the overall signal of epistasis.

#### Robustness to population structure and imperfectly tagged causal SNPs

Population stratification can increase the false positive rate in GWAS and is commonly accounted for by including principal components (PCs) computed from genotype data as covariates in the analysis [30, 31]. A concern is that this approach might not adequately correct for the confounding effects of population stratification on tests of epistasis effects. To explore the effect of population stratification, we reran our analyses on trait-gene pairs previously discovered as significant with the number of PCs included as covariates increased to 40 (from 20). We observe a high correlation in the − log_10_ p-values and in the variance component estimates when using 40 vs 20 PCs (Figure 2a; Spearman correlation *ρ* ≈ 1), indicating that our findings are robust to population stratification.

**Figure 2:**
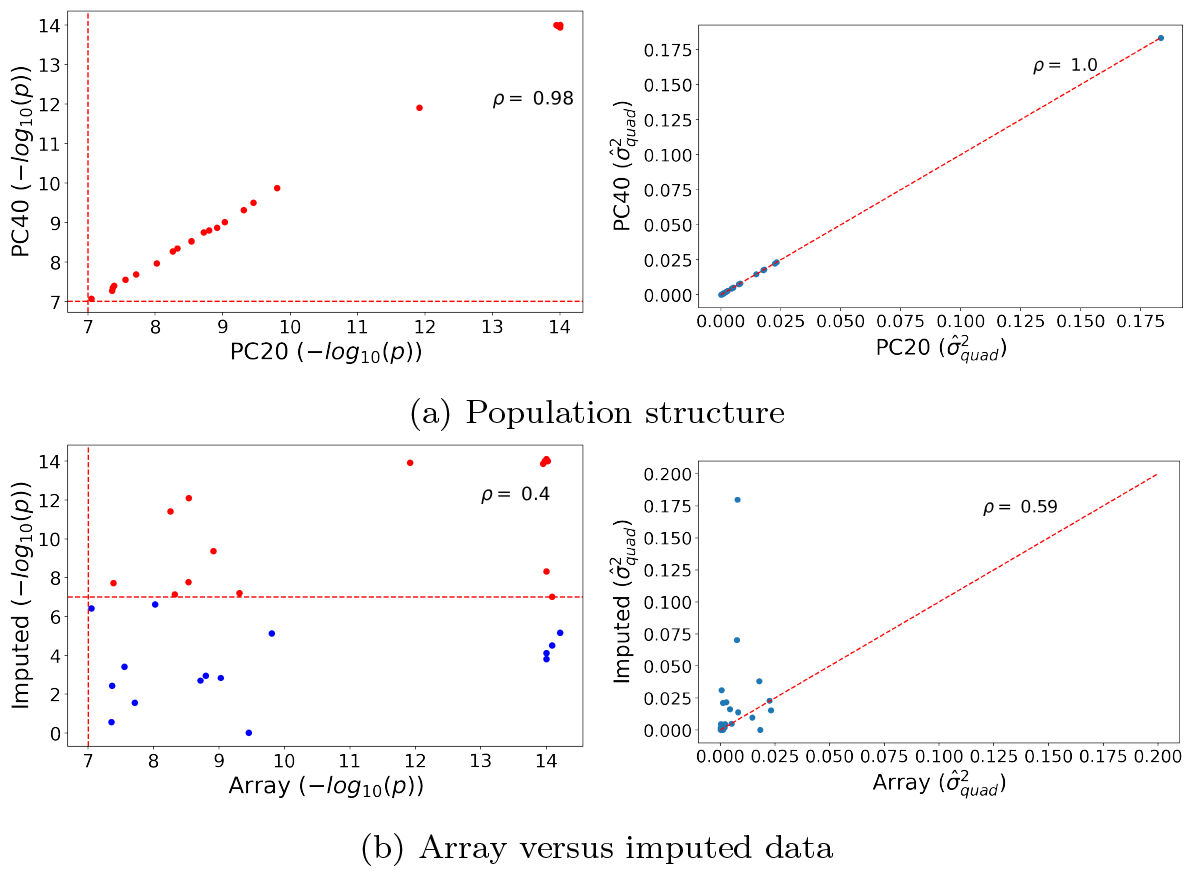
Robustness analysis. We report the Spearman correlation (*ρ*) between estimators of 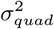 obtained by QuadKAST under different scenarios. (a) We report the correlation of the negative *log*_10_ p-values and the estimates of 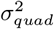 for all the significant epistatic trait-gene pairs when we include the top 20 PCs (default choice) vs the top 40 PCs. (b) We report the correlation of the same estimators between array dataset (20 PC) and imputed dataset. For ease of visualization, estimates than 0.2 have been excluded from the display.

False positive epistatic signals can occur due to imperfect tagging of causal SNPs in the genotyping array [32–34]. To evaluate the robustness of our results in this setting, we simulated a linear additive phenotype from 4,824,392 imputed SNPs (of which 1% SNPs were causal). We then applied QuadKAST to genotypes on the array dataset with SNPs grouped into *filterGenes* protein-coding genes. This experiment simulates a scenario where some of the causal SNPs are missing from the analysis. Figure 3 shows that QuadKAST remains calibrated in this setting (both when we include or exclude self-interactions).

**Figure 3:**
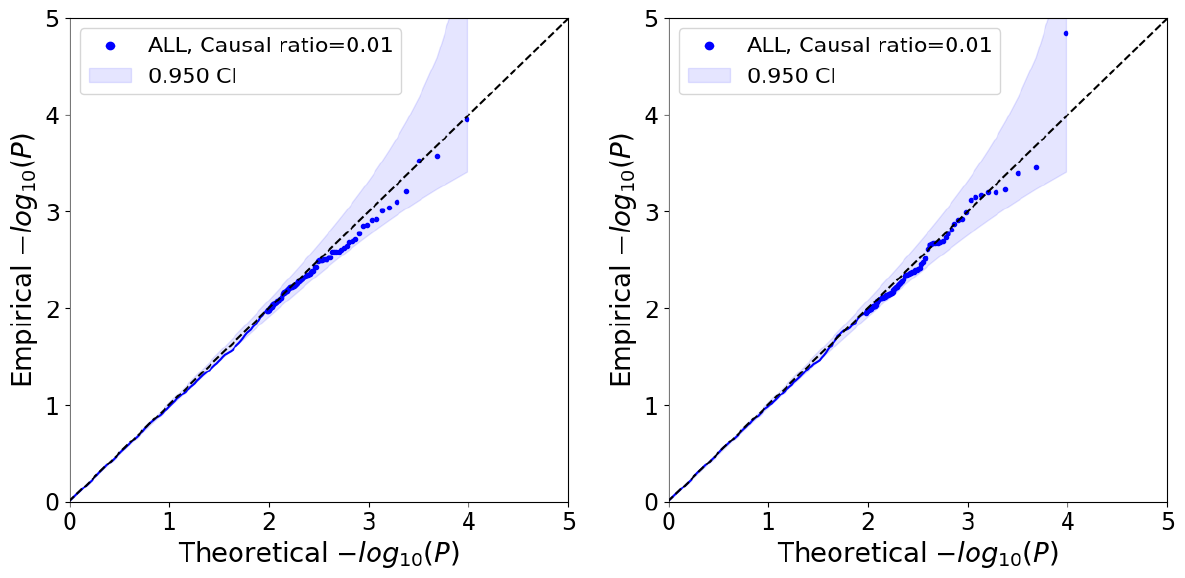
Calibration of QuadKAST when causal variants are imperfectly tagged. We simulated an additive phenotype using imputed genotypes over 4, 824, 392 SNPs (of which we randomly selected 1% of the SNPs to be causal). We then tested QuadKAST on the SNPs genotyped on the UKBB array. On the left, we have the self-interaction results, with *N* = 50K individuals, UK Biobank whole genome data, and gene annotations of up to 50 SNPs, with a total of 9, 515 genes. On the right, we have the no self-interaction results, with the same N, data, and gene annotations.

Beyond validating the calibration of QuadKAST in simulations, we wanted to confirm the consistency of our signals when we include additional SNPs within the sets analyzed. To do this, we re-analyzed the significant trait-gene pairs using the imputed genotype dataset. We observed that 17*/*32 trait-gene pairs were also statistically significant on the imputed genotypes. The Spearman correlation between the − log_10_ p-values on the two SNP sets was 0.4 while the Spearman correlation of the estimated variance components was 0.59 (Figure 2b). A detailed comparison of the estimated variance component across different robustness tests can be found in Supplementary Table S5.

#### Characterizing the importance of individual interactions

To dissect the contributions of individual interactions, we compute the posterior probability distribution of the effects of the interactions (at the MLE or REML estimates of the variance components). The posterior probability of the vector of interaction effects, ***γ***, is described by a multivariate normal distribution. This allows us to derive the posterior mean and variance for each interaction which can then be used to assess the importance of each pair of SNPs in explaining the set-level signal. Given the posterior mean *μ*_*t*_ and standard deviation *σ*_*t*_ of an interaction *t*, we assign an importance score for this interaction as 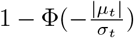 where Φ is the CDF of the standard normal distribution (we caution that these importance scores are not p-values but merely summaries of the posterior distribution of effects). We computed these measures of importance for each interaction at each of 17 epistatic significant gene-trait pairs which passed the significant threshold 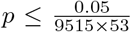 on both array and imputed datasets. Importantly, several trait-gene pairs (Alanine Amnotransferase - PNPLA3 and Mean sphered cell volume - SPTA1) that show significant epistatic effect at the set level do not demonstrate interactions with strong importance scores (significance threshold was defined as 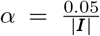, with ***I*** representing the set of interactive features under consideration in one set) (Supplementary Figure S7). This highlight the power of set-based epistatic testing using QuadKAST. One the other hand, genes that show significant epistatic effects while harboring a small number of SNPs typically demonstrated stronger evidence for the presence of individual pairwise interactive effects (Mean sphered cell volume - OR10Z1 and Urate - DNAJC16). We also identify cases where epistasis shows polygenic, strong, and potentially correlated effects (Lipoprotein-A – LPA), in which case further careful analysis is needed to account for potential model misspecification.

## Discussion

We have described QuadKAST, a computationally efficient algorithm that is capable of testing the epistatic effect within a set in an interpretable approach. Applying QuadKAST to 53 quantitative traits in the UK Biobank, we found 32 trait-gene pairs with a strong signal of having epistatic effect. Importantly, the epistatic signal being detected is readily interpretable in the following way. First, the significant epistatic effect detected can be explained by the pairwise interaction model without the intertwining of more complicated interactions. Second, the interactive features and the potentially corresponding weights within the set can be flexibly adjusted to fit specific tasks without compromising the calibration. This flexibility also allows for standardization within the expanded feature space, facilitating the computation of interpretable variance components. Finally, the specific interactive features can be further identified in the downstream analysis, once being detected. Our work lays the groundwork for interpretable, large-scale investigations of epistasis, enabling new insights into its role and significance in complex traits.

Our current work has several limitations. First, the calibration of hypothesis tests and the accuracy of the variance component estimates obtained by QuadKAST may be deteriorated due to inaccuracies in model assumptions, particularly in scenarios where SNP interaction effects exhibit correlated and directional effects. The impact of modeling assumptions on the results from QuadKAST remains an area for further investigation. Second, while QuadKAST enables a deeper understanding of epistasis in local genomic regions such as protein-coding genes investigated in this work, it is important to recognize that interactions within a genomic region represent only a small fraction of the potential forms of epistasis. Third, our set-based strategy requires sets to be defined *a priori*. Alternative approaches to aggregate variants into sets and searching over sets could be combined with the efficient testing within QuadKAST to discover epistasis across the genome. Fourth, the assignment of weights to pairwise interactions as a means of prioritizing specific SNPs or pairs of SNPs can enhance the statistical power and merits further exploration. Finally, it is possible to test for other types of epistasis using QuadKAST including the use of haplotype data although more careful interpretation might be needed to understand the results. We leave these directions for future work.

## 4 Acknowledgments

This research was conducted using the UK Biobank Resource under application 33127. We thank the participants of UK Biobank for making this work possible.

## 5 Code availability

QuadKAST can be found at https://github.com/sriramlab/FastKAST/tree/QuadKAST with the required package installation script, exemplar simulation files, script for running QuadKAST, and results with tutorial analysis. The simulator used in the experiments can be found at https://github.com/alipazokit/simulator. SKAT (v.2.2.5) can be found at https://cran.r-project.org/web/packages/SKAT/index.html.

## Notes

### Competing Interest Statement

The authors have declared no competing interest.

