## Supplementary materials for "A Scalable Adaptive Quadratic Kernel Method for Interpretable Epistasis Analysis in Complex Traits"

### Supplementary Information

#### Supplementary figures

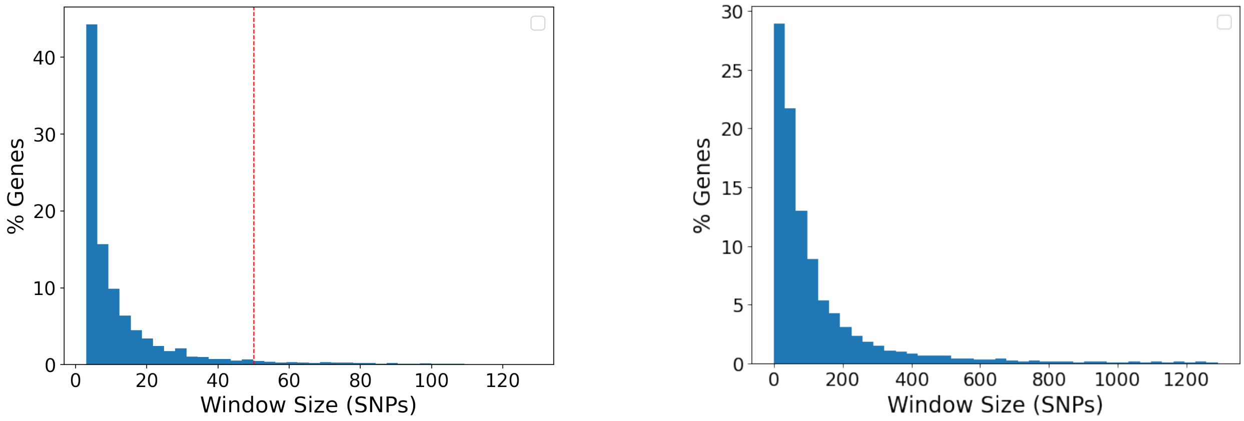


(a) UKBB array data (b) UKBB imputed data

Figure S1: **Distribution of the number of SNPs in protein-coding genes**. (a) The effective SNP number distribution on the array genotype dataset; the red line indicates the 50 SNP cutoff used for real data testing. (b) The effective SNP number distribution on the imputed genotype dataset. In both (a) and (b), we limited the display of gene sizes to the 99th percentile for visualization purposes.

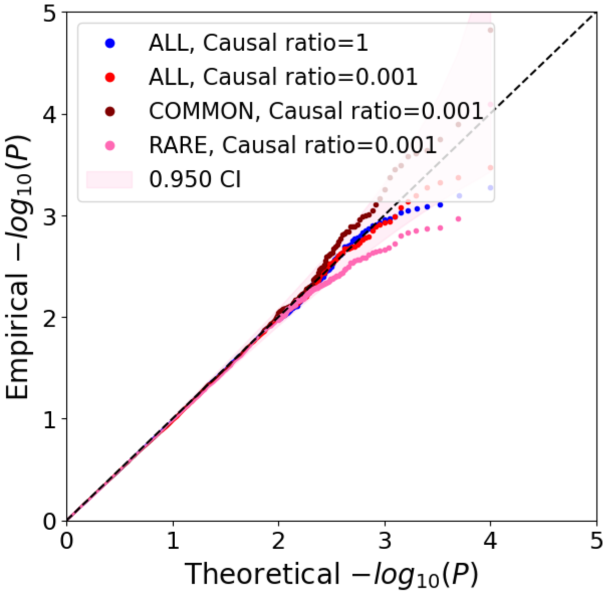


(a) Self-interaction

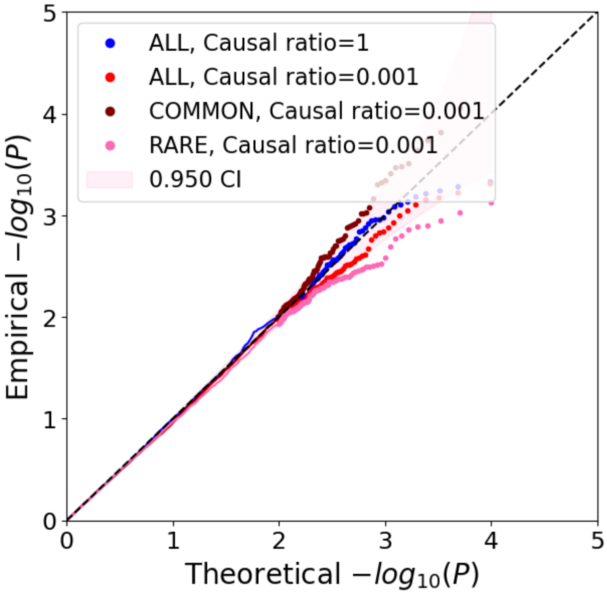


(b) No Self-interaction

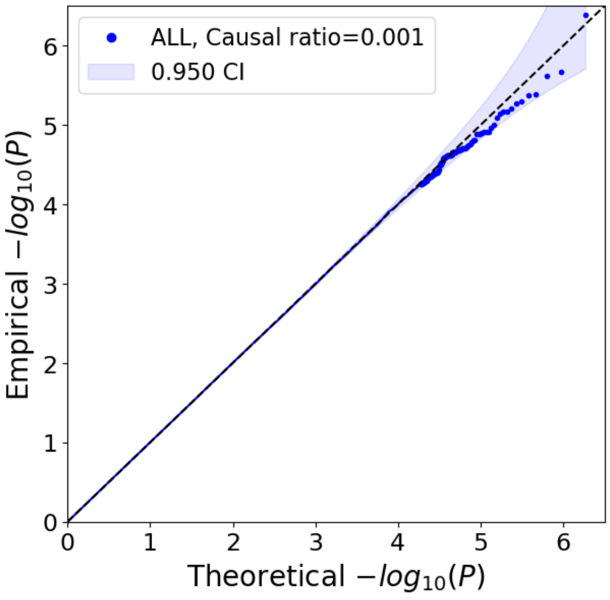


(c) Self-interaction

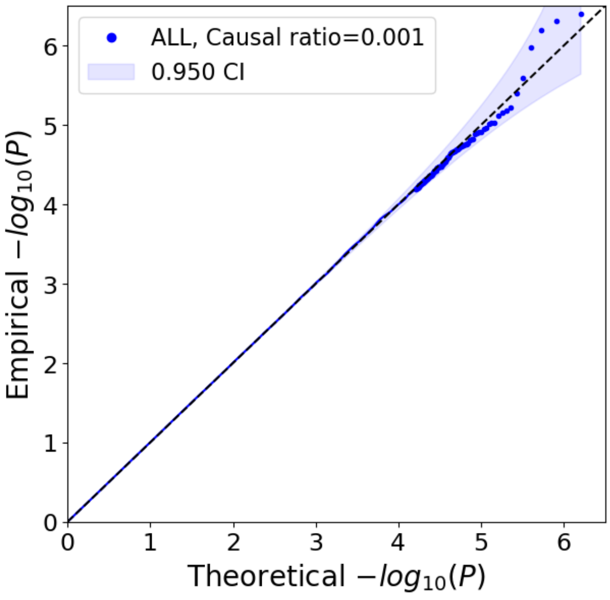


(d) No Self-interaction

Figure S2: **Calibration of QuadKAST.** (a) We simulated under four common genetic architectures with additive effects only (N = 50K individuals, UK Biobank whole genome data). A detailed description of the simulations can be found in the Calibration of QuadKAST section. We then tested the simulations on $9,515$ protein-coding gene annotations (genes with $> 50$ SNPs excluded) from the UK Biobank array dataset under the self-interaction included setting. (b) We simulated under the identical genetic architectures described in (a) while performing the test under the self-interaction excluded setting. (c) We repeated the calibration test on 200 phenotypes simulated purely with linear effects (ALL, Causal ratio = 0.001) to account for higher p-value precision. (d) We performed the exact same simulation as (c) but tested under the self-interaction excluded setting.

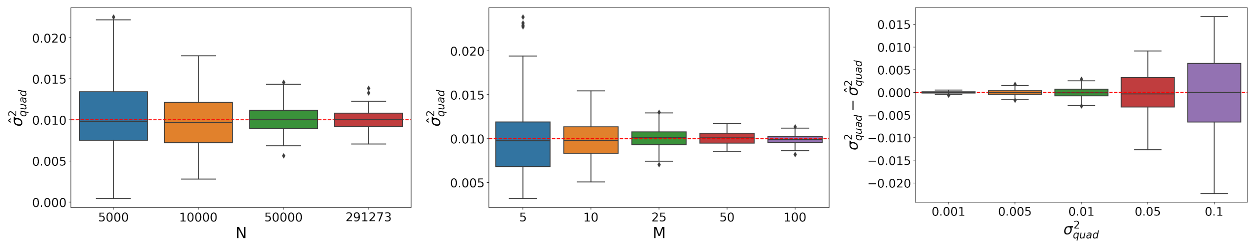


Figure S3: **Variance component estimation on UKBB data**. We vary each of the following parameters: sample size ($N$), feature size ($M$), and signal strength ($\sigma^2_{quad}$), while keeping other quantities fixed. The default values are ($N = 291,273$, $M = 25$, $\sigma^2_{quad} = 0.01$). For each setting, we performed the analysis with 100 replicates.

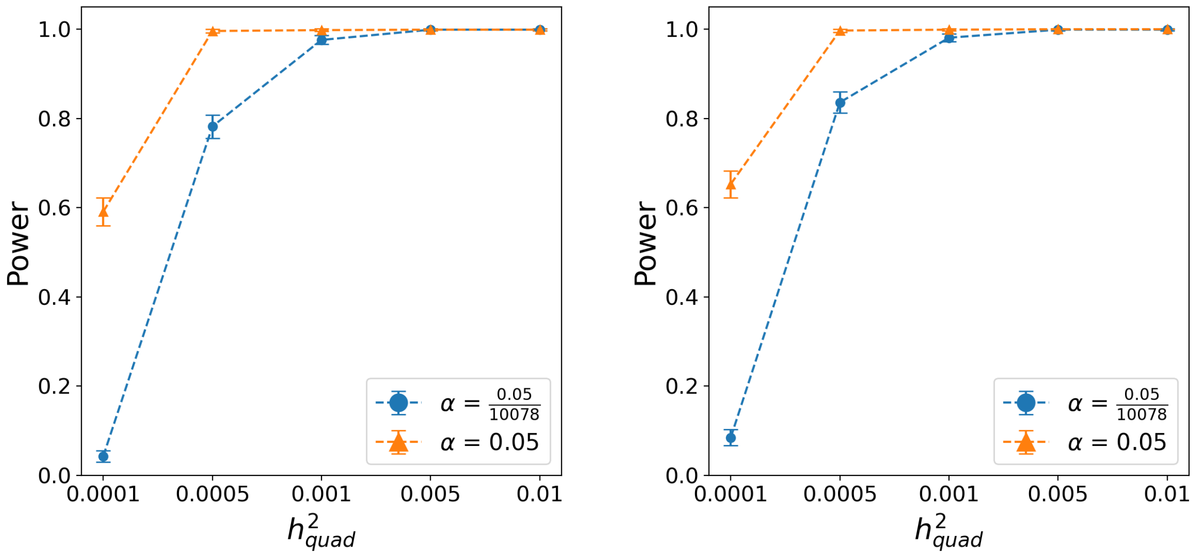


(a) Self-interaction

(b) No self-interaction

Figure S4: **Power analysis of QuadKAST on simulated data.** We randomly selected 5K individuals and 1,000 protein-coding genes from the UKBB to simulate phenotypes with (a) and without self-interaction effects (b). For each set, we varied the heritability (explained completely by the quadratic effect) and computed the expected power of QuadKAST at different thresholds.

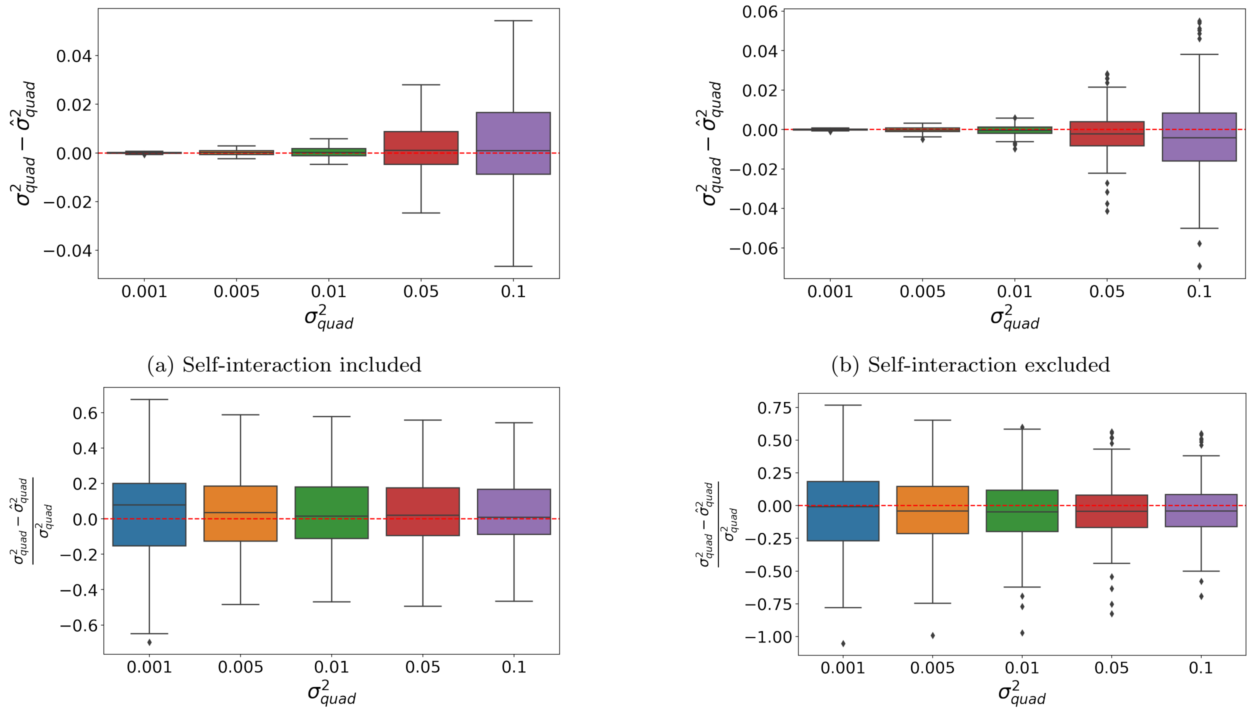


(a) Self-interaction included

(b) Self-interaction excluded

(c) Self-interaction included (relative error)

(d) Self-interaction excluded (relative error)

Figure S5: **Accuracy of variance component estimation** (a) The simulation and estimation of the quadratic component $\sigma^2_{quad}$ are performed by including the self-interaction of the variants in the target set. (b) The simulation and estimation of the quadratic component $\sigma^2_{quad}$ are performed by excluding the self-interaction of the variants in the target set. In order to conduct simulation under the actual SNP distributions, we randomly select 100 genes from the total available genes with outliers (actual SNPs number less than 4 or greater than 50) removed. The sample size used in the simulation was $N = 291,273$. (c) and (d) were done with the exact same experimental settings as (a) and (b), but the relative error $\left(\frac{\sigma^2_{quad} - \hat{\sigma}^2_{quad}}{\sigma^2_{quad}}\right)$ was computed and displayed instead.

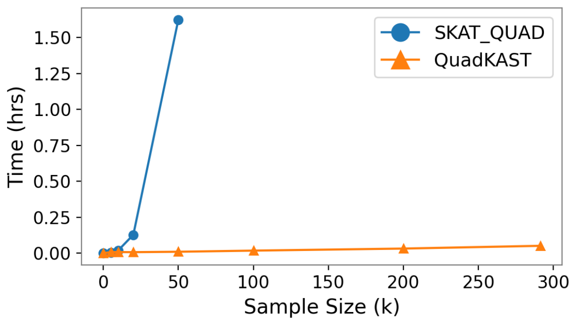


(a) Time

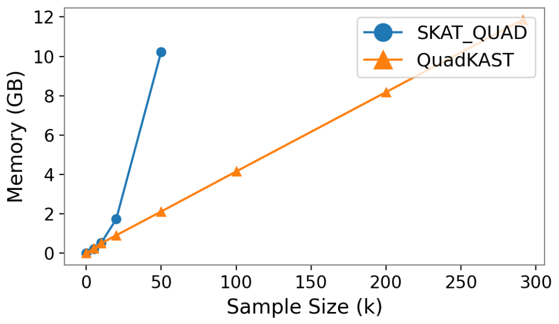


(b) Memory

Figure S6: **Scaling of computational and memory requirements.** Averaged time and memory consumption of quadratic kernel and SVD over ten trials using QuadKAST and SKAT Quadratic (SKAT_QUAD) (varying sample size applied to a set containing 100 SNPs typed on the UKBB array).

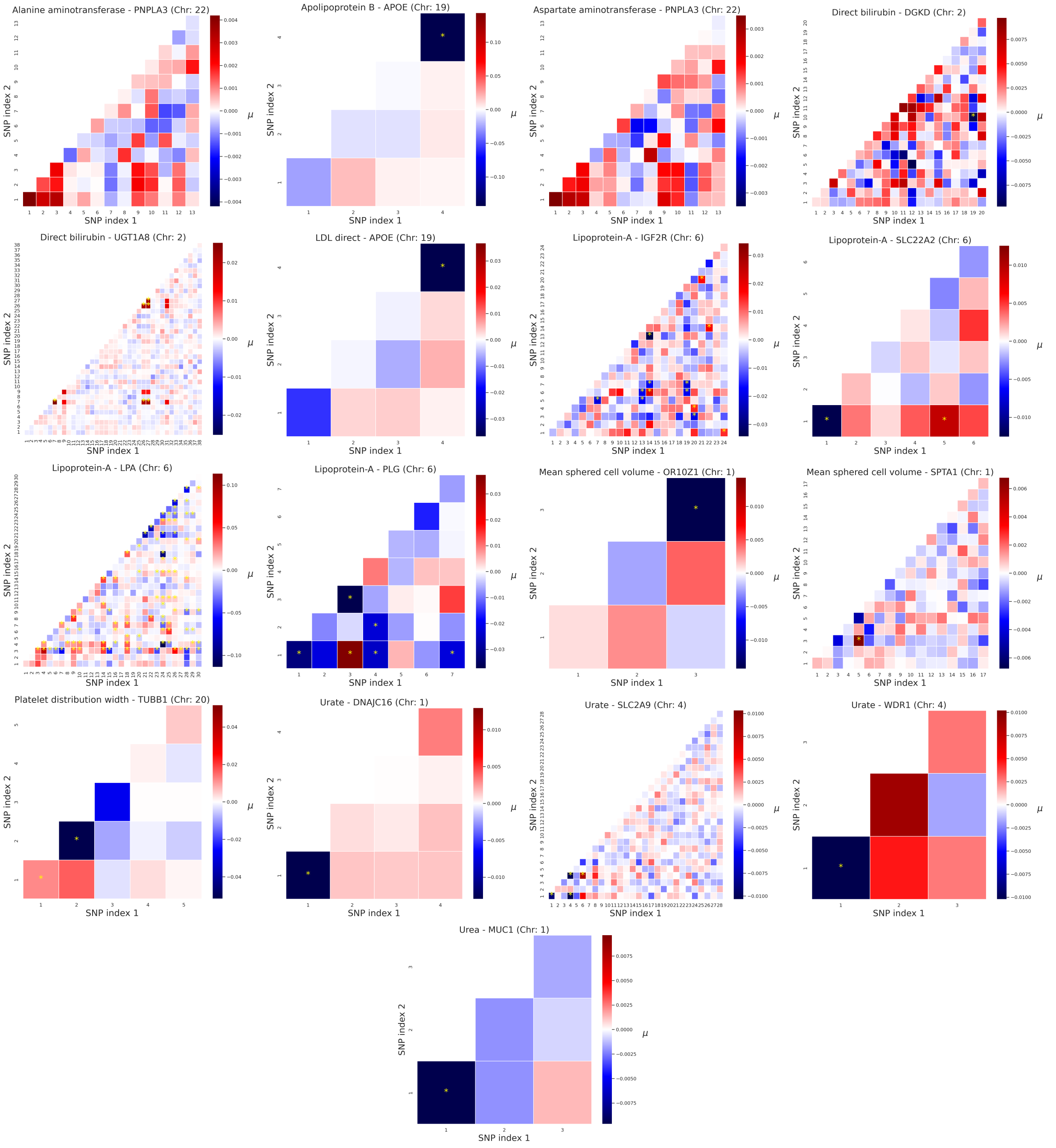


Figure S7: **Analysis of the contribution of each interaction for significant gene-trait pairs**. The axes denotes the SNP index within a set. The color indicates the strength of the interaction as quantified by its posterior mean. The cell with interaction importance score passing the Bonferroni correction threshold $0.05/|\boldsymbol{I}|$ were denoted with yellow star, where $\boldsymbol{I}$ indicates the set of interactive features in a given gene.

#### Supplementary tables

| Category | Trait | Category | Trait |
|---|---|---|---|
| Anthropometry | Body mass index | Glucose metabolism | Hemoglobin A1c |
| | BMD Heel T-score | Kidney | Urea |
| | Height | | Urate |
| | Basal metabolic rate | | Sodium in urine |
| Blood biochemistry | High light scatter reticulocyte count | | Creatinine |
| | White blood cell count | | Creatinine in urine |
| | IGF-1 | | Potassium in urine |
| | RBC distribution width | | Microalbumin in urine |
| | RBC count | | Cystatin-C |
| | Platelet distribution width | Lipid metabolism | Triglycerides |
| | Platelet count | | Cholesterol |
| | Monocyte count | | LDL direct |
| | Eosinophil count | | Apolipoprotein A |
| | C-reactive protein | | HDL cholesterol |
| | Testosterone | Liver | Total bilirubin |
| | Mean corpuscular hemoglobin | | Direct bilirubin |
| | Phosphate | | Aspartate aminotransferase |
| | Mean sphered cell volume | | Alkaline phosphatase |
| | Mean platelet volume | | Alanine aminotransferase |
| | Albumin | Lung | FVC |
| | SHBG | | FEV1-FVC ratio |
| | Lymphocyte count | Other | Corneal Hysteresis |
| Blood pressure | Diastolic blood pressure | | Calcium |
| | Systolic blood pressure | | Alcohol intake frequency |
| Cardiovascular | Lipoprotein-A | | Age first birth |
| | Apolipoprotein B | Renal | Total protein |
| Glucose metabolism | Glucose | | |

Table S1: **Trait category information.** This table presents the trait and the corresponding category information for all the 53 quantitative traits.

| Threshold ($\alpha$) | $10^{-1}$ | $10^{-2}$ | $10^{-3}$ | $10^{-4}$ | $10^{-5}$ | $10^{-6}$ |
|---|---|---|---|---|---|---|
| FPR ($\times\alpha$) | 1.004 | 1.010 | 0.988 | 1.020 | 0.694 | 0.534 |
| p-value | 0.045 | 0.182 | 0.604 | 0.784 | 0.186 | 0.524 |

Table S2: **Calibration analysis** (Self-interaction included). False positive rate for varying p-value thresholds corresponding to Supplementary Figure S2c. After filtering out genes with SNPs number less than three, we obtained 1,872,415 p-values from the simulation setting (ALL, Causal ratio = 0.001) from which we report the FPR. The p-value row describes whether the reported FPR significantly differs from the expected FPR for the given threshold. None of the p-values are significant after correcting for multiple testing suggesting that the FPR is calibrated across significance thresholds.

| Threshold ($\alpha$) | $10^{-1}$ | $10^{-2}$ | $10^{-3}$ | $10^{-4}$ | $10^{-5}$ | $10^{-6}$ |
|---|---|---|---|---|---|---|
| FPR ($\times\alpha$) | 1.005 | 1.008 | 1.004 | 0.979 | 0.806 | 1.859 |
| P-value | 0.026 | 0.287 | 0.878 | 0.790 | 0.435 | 0.275 |

Table S3: **Calibration analysis** (Self-interaction excluded). False positive rate for varying p-value thresholds corresponding to Supplementary Figure S2d. After filtering out genes with SNPs number less than three, we obtained 1,613,833 p-values from the simulation setting (ALL, Causal ratio = 0.001) from which we report the FPR. The p-value row describes whether the reported FPR significantly differs from the expected FPR for the given threshold. None of the p-values are significant after correcting for multiple testing suggesting that the FPR is calibrated across significance thresholds.

Table S4: **Epistatic and additive variance component comparison of the significant trait-gene pairs**. We quantified the variance explained by the additive effect as $\sigma_g^2$, and the variance explained by the quadratic effect after excluding the additive effect as $\sigma_{quad}^2$, and the corresponding standard error (termed $SE(\sigma_g^2)$ and $SE(\sigma_{quad}^2)$).

| Trait | CHR | Gene | Start (MB) | End (MB) | $\sigma_g^2$ $\times 10^{-3}$ | $SE(\sigma_g^2)$ $\times 10^{-3}$ | $\sigma_{quad}^2$ $\times 10^{-3}$ | $SE(\sigma_{quad}^2)$ $\times 10^{-3}$ |
|---|---|---|---|---|---|---|---|---|
| Alanine aminotransferase | 22 | PNPLA3 | 44.32 | 44.34 | 2.72 | 1.26 | 0.34 | 0.11 |
| | 22 | SAMM50 | 44.35 | 44.39 | 2.34 | 1.18 | 0.17 | 0.08 |
| Apolipoprotein B | 19 | BCAM | 45.31 | 45.32 | 8.14 | 4.41 | 2.72 | 1.24 |
| | 19 | APOE | 45.41 | 45.41 | 64.21 | 45.42 | 22.48 | 10.26 |
| Aspartate aminotransferase | 22 | PNPLA3 | 44.32 | 44.34 | 2.73 | 1.26 | 0.34 | 0.11 |
| Creatinine | 1 | DNAJC16 | 15.86 | 15.89 | 0.17 | 0.15 | 0.37 | 0.24 |
| Direct bilirubin | 2 | SAG | 234.22 | 234.26 | 21.99 | 11.02 | 14.68 | 4.34 |
| | 2 | DGKD | 234.26 | 234.38 | 72.58 | 23.24 | 4.49 | 1.02 |
| | 2 | USP40 | 234.39 | 234.47 | 23.30 | 11.28 | 1.12 | 0.35 |
| | 2 | UGT1A8 | 234.53 | 234.68 | 74.98 | 18.98 | 18.18 | 1.87 |
| Eosinophil count | 11 | PRG3 | 57.14 | 57.15 | 0.36 | 0.30 | 8.11 | 4.76 |
| HDL cholesterol | 16 | CETP | 57.00 | 57.02 | 17.64 | 6.99 | 0.19 | 0.08 |
| Hemoglobin A1c | 10 | HK1 | 71.05 | 71.16 | 18.78 | 4.62 | 1.03 | 0.22 |
| LDL direct | 19 | APOE | 45.41 | 45.41 | 35.18 | 24.89 | 1.99 | 1.06 |
| Lipoprotein-A | 6 | IGF2R | 160.39 | 160.53 | 40.00 | 11.65 | 23.22 | 3.27 |
| | 6 | SLC22A2 | 160.64 | 160.68 | 24.68 | 14.27 | 0.53 | 0.24 |
| | 6 | SLC22A3 | 160.77 | 160.87 | 86.38 | 35.30 | 7.61 | 1.66 |
| | 6 | LPA | 160.95 | 161.09 | 546.11 | 141.81 | 183.37 | 14.48 |
| | 6 | PLG | 161.12 | 161.17 | 176.95 | 94.65 | 7.84 | 2.71 |
| | 6 | AGPAT4 | 161.56 | 161.65 | 23.36 | 7.63 | 17.75 | 2.56 |
| Mean corpuscular hemoglobin | 22 | TMPRSS6 | 37.46 | 37.50 | 10.15 | 3.51 | 0.70 | 0.23 |
| Mean sphered cell volume | 1 | OR10Z1 | 158.58 | 158.58 | 7.66 | 6.26 | 0.31 | 0.21 |
| | 1 | SPTA1 | 158.58 | 158.66 | 9.58 | 3.62 | 0.74 | 0.20 |
| Monocyte count | 22 | CECR1 | 17.66 | 17.69 | 0.12 | 0.08 | 0.23 | 0.10 |
| Platelet distribution width | 20 | TUBB1 | 57.59 | 57.60 | 21.44 | 13.83 | 5.20 | 2.52 |
| | 20 | EDN3 | 57.88 | 57.90 | 0.85 | 0.37 | 1.17 | 0.29 |
| SHBG | 17 | TNK1 | 7.29 | 7.29 | 1.53 | 1.50 | 0.28 | 0.20 |
| Urate | 1 | DNAJC16 | 15.86 | 15.89 | 0.25 | 0.21 | 0.38 | 0.26 |
| | 4 | SLC2A9 | 9.83 | 10.03 | 23.76 | 6.38 | 2.17 | 0.28 |
| | 4 | WDR1 | 10.08 | 10.12 | 9.61 | 7.86 | 0.28 | 0.20 |
| | 4 | MEPE | 88.76 | 88.77 | 0.38 | 0.28 | 0.28 | 0.16 |
| Urea | 1 | MUC1 | 155.16 | 155.16 | 0.82 | 0.68 | 0.17 | 0.12 |

Table S5: **Variance components comparison across different settings**. We tested the data with QuadKAST using three settings: default settings, using 40 PCs to control for population structure, and using an imputed dataset. We compared the estimated variance component obtained from each setting.

| Trait | CHR | Gene | Start (MB) | End (MB) | $\sigma^2_{quad}$ $\times 10^{-3}$ | $\sigma^2_{quad}$ (40 PC) $\times 10^{-3}$ | $\sigma^2_{quad}$ (Imputed) $\times 10^{-3}$ |
|---|---|---|---|---|---|---|---|
| Alanine aminotransferase | 22 | PNPLA3 | 44.32 | 44.34 | 0.34 | 0.34 | 0.57 |
| | 22 | SAMM50 | 44.35 | 44.39 | 0.17 | 0.17 | 0.52 |
| Apolipoprotein B | 19 | BCAM | 45.31 | 45.32 | 2.72 | 2.73 | 21.82 |
| | 19 | APOE | 45.41 | 45.41 | 22.48 | 22.41 | 22.98 |
| Aspartate aminotransferase | 22 | PNPLA3 | 44.32 | 44.34 | 0.34 | 0.34 | 0.48 |
| Creatinine | 1 | DNAJC16 | 15.86 | 15.89 | 0.37 | 0.37 | 0.14 |
| Direct bilirubin | 2 | SAG | 234.22 | 234.26 | 14.68 | 14.69 | 9.68 |
| | 2 | DGKD | 234.26 | 234.38 | 4.49 | 4.51 | 16.43 |
| | 2 | USP40 | 234.39 | 234.47 | 1.12 | 1.13 | 21.25 |
| | 2 | UGT1A8 | 234.53 | 234.68 | 18.18 | 18.16 | 0.00 |
| Eosinophil count | 11 | PRG3 | 57.14 | 57.15 | 8.11 | 8.11 | 13.98 |
| HDL cholesterol | 16 | CETP | 57.00 | 57.02 | 0.19 | 0.19 | 0.30 |
| Hemoglobin A1c | 10 | HK1 | 71.05 | 71.16 | 1.03 | 1.03 | 0.55 |
| LDL direct | 19 | APOE | 45.41 | 45.41 | 1.99 | 1.98 | 2.10 |
| Lipoprotein-A | 6 | IGF2R | 160.39 | 160.53 | 23.22 | 23.20 | 15.45 |
| | 6 | SLC22A2 | 160.64 | 160.68 | 0.53 | 0.54 | 31.10 |
| | 6 | SLC22A3 | 160.77 | 160.87 | 7.61 | 7.62 | 70.18 |
| | 6 | LPA | 160.95 | 161.09 | 183.37 | 183.26 | 792.13 |
| | 6 | PLG | 161.12 | 161.17 | 7.84 | 7.86 | 179.78 |
| | 6 | AGPAT4 | 161.56 | 161.65 | 17.75 | 17.77 | 38.29 |
| Mean corpuscular hemoglobin | 22 | TMPRSS6 | 37.46 | 37.50 | 0.70 | 0.70 | 0.34 |
| Mean sphered cell volume | 1 | OR10Z1 | 158.58 | 158.58 | 0.31 | 0.31 | 0.32 |
| | 1 | SPTA1 | 158.58 | 158.66 | 0.74 | 0.74 | 0.89 |
| Monocyte count | 22 | CECR1 | 17.66 | 17.69 | 0.23 | 0.23 | 1.71 |
| Platelet distribution width | 20 | TUBB1 | 57.59 | 57.60 | 5.20 | 5.21 | 4.91 |
| | 20 | EDN3 | 57.88 | 57.90 | 1.17 | 1.17 | 0.00 |
| SHBG | 17 | TNK1 | 7.29 | 7.29 | 0.28 | 0.28 | 1.04 |
| Urate | 1 | DNAJC16 | 15.86 | 15.89 | 0.38 | 0.37 | 0.23 |
| | 4 | SLC2A9 | 9.83 | 10.03 | 2.17 | 2.18 | 4.77 |
| | 4 | WDR1 | 10.08 | 10.12 | 0.28 | 0.28 | 4.77 |
| | 4 | MEPE | 88.76 | 88.77 | 0.28 | 0.28 | 0.05 |
| Urea | 1 | MUC1 | 155.16 | 155.16 | 0.17 | 0.17 | 0.24 |

### S1 Supplementary Notes

#### S1.1 Score statistics

Under the assumption that $\boldsymbol{y} \sim \mathcal{N}(\boldsymbol{X\alpha}, \sigma^2_{quad}\boldsymbol{K} + \sigma^2_\epsilon \boldsymbol{I})$, we can write down the log-likelihood as

$$\ell(\boldsymbol{\alpha}, \sigma^2_{quad}, \sigma^2_\epsilon) = -\frac{1}{2}[\log|\sigma^2_{quad}\boldsymbol{K} + \sigma^2_\epsilon \boldsymbol{I}| + (\boldsymbol{y} - \boldsymbol{X\alpha})^T(\sigma^2_{quad}\boldsymbol{K} + \sigma^2_\epsilon \boldsymbol{I})^{-1}(\boldsymbol{y} - \boldsymbol{X\alpha})] + Const$$

Under the null hypothesis $\sigma^2_{quad} = 0$, the constrained MLE of $(\boldsymbol{\alpha}, \sigma^2_\epsilon)$:

$$\hat{\boldsymbol{\alpha}} = (\boldsymbol{X^T X})^{-1}\boldsymbol{X}^T\boldsymbol{y}$$
$$\hat{\sigma}^2_\epsilon = \frac{\boldsymbol{y}^T\boldsymbol{Py}}{N - K}$$

We can further write down the partial derivative of $\ell$ with respect to $\sigma^2_{quad}$ as

$$\frac{\partial \ell(\boldsymbol{\alpha}, \sigma^2_{quad}, \sigma^2_\epsilon)}{\sigma^2_{quad}} = \frac{1}{2}(\boldsymbol{y} - \boldsymbol{X\alpha})^T\boldsymbol{\Sigma}^{-1}\boldsymbol{K}\boldsymbol{\Sigma}^{-1}(\boldsymbol{y} - \boldsymbol{X\alpha}) - \frac{1}{2}tr(\boldsymbol{\Sigma}^{-1}\boldsymbol{K})$$

where $\boldsymbol{\Sigma} = \sigma^2_{quad}\boldsymbol{K} + \sigma^2_\epsilon \boldsymbol{I}$.
Plugging in the parameter estimates under the null hypothesis:

$$\begin{aligned}
\left.\frac{\partial \ell(\hat{\boldsymbol{\alpha}}, \sigma^2_{quad}, \hat{\sigma}^2_\epsilon)}{\partial \sigma^2_{quad}}\right|_{\sigma^2_{quad}=0} &= \frac{1}{2\hat{\sigma}^4_\epsilon}(\boldsymbol{y} - \boldsymbol{X}\hat{\boldsymbol{\alpha}})^T\boldsymbol{K}(\boldsymbol{y} - \boldsymbol{X}\hat{\boldsymbol{\alpha}}) - \frac{1}{2\hat{\sigma}^2_\epsilon}tr(\boldsymbol{K}) \\
&= \frac{1}{2\hat{\sigma}^4_\epsilon}(\boldsymbol{y} - \boldsymbol{X}(\boldsymbol{X^T X})^{-1}\boldsymbol{X}^T\boldsymbol{y})^T\boldsymbol{K}(\boldsymbol{y} - \boldsymbol{X}(\boldsymbol{X^T X})^{-1}\boldsymbol{X}^T\boldsymbol{y}) - \frac{1}{2\hat{\sigma}^2_\epsilon}tr(\boldsymbol{K}) \\
&= \frac{1}{2\hat{\sigma}^4_\epsilon}\boldsymbol{y}^T\boldsymbol{PKPy} - \frac{1}{2\hat{\sigma}^2_\epsilon}tr(\boldsymbol{K})
\end{aligned}$$

This leads to the score test statistic that is used in SKAT [1]: $Q = \frac{1}{\hat{\sigma}^2_\epsilon}\boldsymbol{y}^T\boldsymbol{PKPy}$. An alternative score statistic for which the exact sampling distribution under the null hypothesis can be computed has been proposed in [35].

##### S1.1.1 Sampling distribution of the score statistic

For completeness, we derive the sampling distribution of the score statistic, following closely the treatment in [35].

Let $\mathcal{C}()$ denote the column space of a matrix. Let $R$ and $S$ denote the ranks of $\boldsymbol{X}$ and $\boldsymbol{PKP}$ respectively. Let $\boldsymbol{A}_0 \in \mathbb{R}^{N\times R}$ denote an orthonormal basis of $\mathcal{C}(\boldsymbol{X})$ and $\boldsymbol{A}_1 \in \mathbb{R}^{N\times S}$ denote an orthonormal basis based on an eigendecomposition of $\boldsymbol{PKP}$. $\boldsymbol{A}_2 \in \mathbb{R}^{N\times(N-R-S)}$ is an orthonormal basis of $\mathcal{C}(\boldsymbol{A}_0, \boldsymbol{A}_1)^\perp = \mathcal{C}(\boldsymbol{X}, \boldsymbol{A}_1)^\perp$ so that $\boldsymbol{A} = [\boldsymbol{A}_1, \boldsymbol{A}_2] \in \mathbb{R}^{N\times(N-R)}$ is an orthonormal basis of $\mathcal{C}(\boldsymbol{X})^\perp$.
Using the uniqueness of orthogonal projections, we have $\boldsymbol{P} = \boldsymbol{AA}^{\mathrm{T}}$. Under the null hypothesis:

$$\begin{aligned}
\boldsymbol{w} &:= \boldsymbol{A}^{\mathrm{T}}\boldsymbol{y} \\
&\sim \mathcal{N}(\boldsymbol{A}^{\mathrm{T}}\boldsymbol{X\alpha}, \boldsymbol{A}^{\mathrm{T}}\boldsymbol{I}_N\sigma^2_\epsilon\boldsymbol{A}) \\
&\sim \mathcal{N}(\boldsymbol{0}, \boldsymbol{I}_{N-R}\sigma^2_\epsilon) \qquad (7)
\end{aligned}$$

We also have:

$$\begin{aligned}
\boldsymbol{PKP} &= \boldsymbol{A}_1 diag(\lambda_1, \ldots, \lambda_S)\boldsymbol{A}_1^{\mathrm{T}} \\
&= [\boldsymbol{A}_1, \boldsymbol{A}_2] diag(\lambda_1, \ldots, \lambda_S, 0, \ldots, 0)[\boldsymbol{A}_1, \boldsymbol{A}_2]^{\mathrm{T}} \\
&= \boldsymbol{A} diag(\lambda_1, \ldots, \lambda_S, 0, \ldots, 0)\boldsymbol{A}^{\mathrm{T}} \qquad (8)
\end{aligned}$$

Here $(\lambda_1, \ldots, \lambda_S)$ are the non-zero eigenvalues of $\boldsymbol{PKP}$.

Combining Equations 7 and 8:

$$\begin{aligned} \boldsymbol{y}^{\mathrm{T}}\boldsymbol{PKPy} &= \boldsymbol{y}^{\mathrm{T}}\boldsymbol{A}diag(\lambda_1,\dots,\lambda_S,0,\dots,0)\boldsymbol{A}^{\mathrm{T}}\boldsymbol{y} \\ &= \boldsymbol{w}^{\mathrm{T}}diag(\lambda_1,\dots,\lambda_S,0,\dots,0)\boldsymbol{w} \\ &= \sigma_\epsilon^2\sum_{i=1}^{S}\lambda_i\chi_i^2 \end{aligned} \tag{9}$$

Here $\chi_i^2$ are independent random variables with a $\chi^2$, 1-df distribution.
Since $\hat{\sigma}_\epsilon^2 \xrightarrow{p} \sigma_\epsilon^2$, we then have

$$Q \xrightarrow{d} \sum_{i=1}^{S}\lambda_i\chi_i^2 \tag{10}$$

##### S1.1.2 Computing the p-value of the score statistic

To compute the score statistic, we write the statistic as: $Q = \frac{1}{D\hat{\sigma}_\epsilon^2}\boldsymbol{y}^{\mathrm{T}}\boldsymbol{P\Phi\Phi}^{\mathrm{T}}\boldsymbol{Py}$. The statistic can be computed in $\mathcal{O}(ND)$ time. Computing the p-value using the sampling distribution in Equation 9 requires computing the eigenvalues of $\boldsymbol{PKP}$. The eigenvalues are obtained as the squared singular values of $\boldsymbol{P\Phi} = \boldsymbol{\Phi} - \boldsymbol{X}\left(\boldsymbol{X}^T\boldsymbol{X}\right)^{-1}\boldsymbol{X}^T\boldsymbol{\Phi}$. The matrix $\boldsymbol{P\Phi}$ can be formed in $\mathcal{O}(NKD + NK^2 + K^3)$ times and its singular values can be computed in $\mathcal{O}(ND^2)$ time (for $D < N$).

#### S1.2 Variance components estimation

When no covariates are included, we have $\boldsymbol{y} \sim \mathcal{N}(0, \sigma^2_{quad}\boldsymbol{K} + \sigma^2_\epsilon \boldsymbol{I})$. We can estimate $(\sigma^2_{quad}, \sigma^2_\epsilon)$ by maximizing the log-likelihood:

$$\ell(\sigma^2_{quad}, \sigma^2_\epsilon) = -\frac{1}{2}\left[\log|\sigma^2_{quad}\boldsymbol{K} + \sigma^2_\epsilon \boldsymbol{I}| + \boldsymbol{y}^T(\sigma^2_{quad}\boldsymbol{K} + \sigma^2_\epsilon \boldsymbol{I})^{-1}\boldsymbol{y}\right] + Const$$

Defining $\delta = \frac{\sigma^2_\epsilon}{\sigma^2_{quad}}$, the log-likelihood can be re-parameterized as:

$$\ell(\sigma^2_{quad}, \delta) = -\frac{1}{2}\left[\log|\sigma^2_{quad}(\boldsymbol{K} + \delta\boldsymbol{I})| + \boldsymbol{y}^T(\sigma^2_{quad}(\boldsymbol{K} + \delta\boldsymbol{I}))^{-1}\boldsymbol{y}\right] + Const$$

Computing the partial derivative w.r.t. $\sigma^2_{quad}$, we get

$$\frac{\partial \ell(\sigma^2_{quad}, \delta)}{\partial \sigma^2_{quad}} = -\frac{1}{2}[\frac{N}{\sigma^2_{quad}} - \frac{\boldsymbol{y}^T(\boldsymbol{K} + \delta\boldsymbol{I})^{-1}\boldsymbol{y}}{\sigma^4_{quad}}] = 0$$
$$\sigma^2_{quad} = \frac{\boldsymbol{y}^T(\boldsymbol{K} + \delta\boldsymbol{I})^{-1}\boldsymbol{y}}{N}$$

Substituting $\sigma^2_{quad}$ into the log-likelihood function, we get the profile log-likelihood:

$$\ell_P(\delta) = -\frac{1}{2}\left[N\log\frac{1}{N}\boldsymbol{y}^T(\boldsymbol{K} + \delta\boldsymbol{I})^{-1}\boldsymbol{y} + \log|(\boldsymbol{K} + \delta\boldsymbol{I})|\right] + Const \tag{11}$$

Given the eigen-decomposition of $\boldsymbol{K} = \boldsymbol{U}diag(\rho_1, \ldots, \rho_N)\boldsymbol{U}^T$ where $\boldsymbol{U} \in \mathbb{R}^{N\times N}$ is the matrix of eigenvectors and $(\rho_1, \ldots, \rho_N)$ are the eigenvalues of $\boldsymbol{K}$, Equation 11 can be rewritten as

$$\begin{aligned}
\ell_P(\delta) &= -\frac{1}{2}\left[N\log\left(\frac{1}{N}\boldsymbol{y}^T(\boldsymbol{USU}^T + \delta\boldsymbol{I})^{-1}\boldsymbol{y}\right) + \log|(\boldsymbol{USU}^T + \delta\boldsymbol{I})|\right] + Const \\
&= -\frac{1}{2}\left[N\log\left(\frac{1}{N}\boldsymbol{y}^T(\boldsymbol{U}(\boldsymbol{S} + \delta\boldsymbol{I})\boldsymbol{U}^T)^{-1}\boldsymbol{y}\right) + \log|\boldsymbol{U}(\boldsymbol{S} + \delta\boldsymbol{I})\boldsymbol{U}^T|\right] + Const \\
&= -\frac{1}{2}\left[N\log\left(\frac{1}{N}\left(\sum_{i=1}^{N}\frac{\tilde{y}_i^2}{\rho_i + \delta}\right)\right) + \sum_{i=1}^{N}\log(s_i + \delta)\right] + Const \qquad (12)
\end{aligned}$$

Here $\tilde{y}_i$ is the $i^{th}$ entry of $\tilde{\boldsymbol{y}} = \boldsymbol{U}^T\boldsymbol{y}$. Evaluating this function takes $\mathcal{O}(N)$ complexity at each iteration of the optimization algorithm.

The above derivation assumes that $\boldsymbol{K}$ is a full rank matrix. However, the rank of $\boldsymbol{K}$ can often be substantially smaller than $N$ leading to additional efficiency in evaluating an optimizing $\ell_P$. In our application, $\boldsymbol{K} = \frac{\boldsymbol{\Phi\Phi}^T}{D}$ where $\boldsymbol{\Phi} \in \mathbb{R}^{N\times D}$ so that the rank of $\boldsymbol{K}$: $R \leq min(D, N) = D$ when $D < N$. In this setting, we have $\rho_i = 0$ for $i > R$ and we can write $\boldsymbol{U} = [\boldsymbol{U}_1, \boldsymbol{U}_2]$, where $\boldsymbol{U}_1 \in \mathbb{R}^{N\times R}$ and $\boldsymbol{U}_2 \in \mathbb{R}^{N\times(N-R)}$ so that:

$$\tilde{y}_i = \begin{cases} [\boldsymbol{U}_1^T\boldsymbol{y}]_i, i \leq R \\ [\boldsymbol{U}_2^T\boldsymbol{y}]_i, i > R \end{cases}$$

We then can rewrite the profile log-likelihood as:

$$
\begin{aligned}
\ell_P(\delta) &= -\frac{1}{2}\left[N\log\left(\frac{1}{N}\left(\sum_{i=1}^{N}\frac{\tilde{y}_i^2}{\rho_i+\delta}\right)\right)+\sum_{i=1}^{N}\log(\rho_i+\delta)\right]+Const\\
&= -\frac{1}{2}\left[N\log\left(\frac{1}{N}\left(\sum_{i=1}^{R}\frac{\tilde{y}_i^2}{\rho_i+\delta}+\sum_{i=R+1}^{N}\frac{\tilde{y}_i^2}{\delta}\right)\right)+\sum_{i=1}^{P}\log(\rho_i+\delta)+(N-R)\log(\delta)\right]+Const\\
&= -\frac{1}{2}\left[N\log\left(\frac{1}{N}\left(\sum_{i=1}^{R}\frac{\tilde{y}_i^2}{\rho_i+\delta}+\frac{1}{\delta}(\sum_{i=1}^{N}\tilde{y}_i^2-\sum_{i=1}^{R}\tilde{y}_i^2)\right)\right)+\sum_{i=1}^{P}\log(\rho_i+\delta)+(N-R)\log(\delta)\right]+Const\\
&= -\frac{1}{2}\left[N\log\left(\frac{1}{N}\left(-\sum_{i=1}^{R}\frac{\rho_i\tilde{y}_i^2}{(\rho_i+\delta)\delta}+\frac{1}{\delta}\sum_{i=1}^{N}\tilde{y}_i^2\right)\right)+\sum_{i=1}^{P}\log(\rho_i+\delta)+(N-R)\log(\delta)\right]+Const\\
&= -\frac{1}{2}\left[N\log\left(\frac{1}{N}\left(-\sum_{i=1}^{R}\frac{\rho_i\tilde{y}_i^2}{(\rho_i+\delta)\delta}+\frac{1}{\delta}\|\tilde{\boldsymbol{y}}\|_2^2\right)\right)+\sum_{i=1}^{P}\log(\rho_i+\delta)+(N-R)\log(\delta)\right]+Const\\
&= -\frac{1}{2}\left[N\log\left(\frac{1}{N}\left(-\sum_{i=1}^{R}\frac{\rho_i\tilde{y}_i^2}{(\rho_i+\delta)\delta}+\frac{1}{\delta}\|\boldsymbol{y}\|_2^2\right)\right)+\sum_{i=1}^{P}\log(\rho_i+\delta)+(N-R)\log(\delta)\right]+Const
\end{aligned}
$$

Evaluating $\ell_P$ in this setting requires computing $\tilde{y}_i$ and $\rho_i$, $i \leq R$ which can be obtained by a one-time computation of the $R$ non-zero eigenvalues and corresponding eigenvectors of $\boldsymbol{K}$. Computation of these eigenvalues and eigenvectors can be obtained in $\mathcal{O}(ND^2)$ time from a SVD of $\boldsymbol{\Phi}$) while subsequent evaluation of $\ell_P$ requires $\mathcal{O}(D)$ time.

#### S1.3 Variance component estimation with covariates included

When covariates are included in the model, we have: $\boldsymbol{y} \sim \mathcal{N}(\boldsymbol{X}\boldsymbol{\alpha}, \sigma^2_{quad}\boldsymbol{K} + \sigma^2_\epsilon \boldsymbol{I})$. Let $L$ and $S$ denote the ranks of $\boldsymbol{X}$ and $\boldsymbol{PKP}$ respectively. Let the full SVD of $\boldsymbol{X} = \boldsymbol{B}_0\boldsymbol{\Sigma}\boldsymbol{C}_0^T$ where $\boldsymbol{B}_0 \in \mathbb{R}^{N\times N}$, $\boldsymbol{C}_0 \in \mathbb{R}^{K\times K}$, and $\boldsymbol{\Sigma}$ is an $N \times K$ rectangular diagonal matrix such that

$$\boldsymbol{\Sigma}_{ij} = \begin{cases} s_i & \text{If } i = j \text{ and } i \leq L \\ 0 & \text{Otherwise} \end{cases}$$

Let $\boldsymbol{B}_0 = [\boldsymbol{B}_1, \boldsymbol{B}_2]$ where $\boldsymbol{B}_1 \in \mathbb{R}^{N\times L}$ and $\boldsymbol{B}_2 \in \mathbb{R}^{N\times(N-L)}$. Similarly, we define $\boldsymbol{C}_1 \in \mathbb{R}^{K\times L}$ and $\boldsymbol{C}_2 \in \mathbb{R}^{K\times(K-L)}$. We then have $\boldsymbol{X} = \boldsymbol{B}_1 diag(s_1, .., s_R)\boldsymbol{C}_1^T$. Together, we have

$$\boldsymbol{P} = \boldsymbol{I}_N - \boldsymbol{X}(\boldsymbol{X}^T\boldsymbol{X})^{-1}\boldsymbol{X}^T = \boldsymbol{B}_0\boldsymbol{B}_0^T - \boldsymbol{B}_1\boldsymbol{B}_1^T = \boldsymbol{B}_2\boldsymbol{B}_2^T$$

where $\boldsymbol{B}_2$ denotes an orthonormal basis of $\mathcal{C}(\boldsymbol{X})^\perp$ and $\mathcal{C}(\boldsymbol{X})$ is the column space of $\boldsymbol{X}$.

We then define:

$$\begin{aligned} \boldsymbol{\omega} &:= \boldsymbol{B}_2^T\boldsymbol{y} \\ &\sim \mathcal{N}(\boldsymbol{B}_2^T\boldsymbol{X}\boldsymbol{\alpha}, \sigma^2_{quad}\boldsymbol{B}_2^T\boldsymbol{K}\boldsymbol{B}_2 + \sigma^2_\epsilon\boldsymbol{B}_2^T\boldsymbol{I}_N\boldsymbol{B}_2) \\ &\sim \mathcal{N}(\boldsymbol{0}_{N-L}, \sigma^2_{quad}\boldsymbol{B}_2^T\boldsymbol{K}\boldsymbol{B}_2 + \sigma^2_\epsilon\boldsymbol{I}_{N-L}) \end{aligned}$$

Let the eigendecomposition of $\boldsymbol{B}_2^T\boldsymbol{K}\boldsymbol{B}_2 = \boldsymbol{U}_b diag(s_1, \ldots, s_{N-L})\boldsymbol{U}_b^T$ where $\boldsymbol{U}_b$ is a $(N-L)\times(N-L)$ orthonormal matrix of eigenvectors of $\boldsymbol{B}_2^T\boldsymbol{K}\boldsymbol{B}_2$. We then define

$$\begin{aligned} \tilde{\boldsymbol{y}} &:= \boldsymbol{U}_b^T\boldsymbol{\omega} = \boldsymbol{U}_b^T\boldsymbol{B}_2^T\boldsymbol{y} \\ &\sim \mathcal{N}(\boldsymbol{0}_{N-L}, \sigma^2_{quad} diag(s_1, \ldots, s_{N-L}) + \sigma^2_\epsilon\boldsymbol{I}_{N-L}) \end{aligned} \tag{13}$$

From Equation 13, the restricted log-likelihood function can be written as:

$$\ell_R(\sigma^2_{quad}, \sigma^2_\epsilon) = -\frac{N-L}{2}\log(2\pi) - \frac{1}{2}\sum_{i=1}^{N-L}\log\left(s_i\sigma^2_{quad} + \sigma^2_\epsilon\right) - \frac{1}{2}\sum_{i=1}^{N-L}\frac{\tilde{y}_i^2}{s_i\sigma^2_{quad} + \sigma^2_\epsilon} \tag{14}$$

which, in turn, yields the profile restricted log-likelihood (as a function of $\delta = \frac{\sigma^2_\epsilon}{\sigma^2_{quad}}$):

$$\ell_{PR}(\delta) = -\frac{1}{2}[(N-L)\log\frac{1}{N-L}\left(\sum_{i=1}^{N-L}\frac{\tilde{y}_i^2}{s_i+\delta}\right) + \sum_{i=1}^{N-L}\log(s_i+\delta)] + Const \tag{15}$$

The values of $\sigma^2_{quad}$ and $\sigma^2_\epsilon$ that maximize the restricted log-likelihood can be efficiently computed using the same technique as described in Supplementary Note S1.2 once the eigendecomposition of $\boldsymbol{B}_2^{\mathrm{T}}\boldsymbol{K}\boldsymbol{B}_2$ has been computed.

##### S1.3.1 Efficient computation

Using the fact that we have $\boldsymbol{P} = \boldsymbol{B}_2\boldsymbol{B}_2^T$, we write

$$\begin{aligned} \boldsymbol{PKP} &= \boldsymbol{B}_2\boldsymbol{B}_2^T\boldsymbol{K}\boldsymbol{B}_2\boldsymbol{B}_2^T \\ &= \boldsymbol{B}_2\boldsymbol{U}_b diag(s_1, \ldots, s_{N-L})\boldsymbol{U}_b^T\boldsymbol{B}_2^T \\ &= \boldsymbol{B} diag(s_1, \ldots, s_{N-L})\boldsymbol{B}^T \end{aligned}$$

Where $\boldsymbol{B} = \boldsymbol{B}_2\boldsymbol{U}_b$ is a $N \times (N-L)$ matrix with orthonormal columns

$$\begin{aligned} \boldsymbol{B}^T\boldsymbol{B} &= \boldsymbol{U}_b^T\boldsymbol{B}_2^T\boldsymbol{B}_2\boldsymbol{U}_b \\ &= \boldsymbol{U}_b\boldsymbol{I}_{N-L}\boldsymbol{U}_b^T = \boldsymbol{U}_b\boldsymbol{U}_b^T = \boldsymbol{I}_{N-L} \end{aligned}$$

Thus, $\boldsymbol{B}$ is the matrix of eigenvectors of $\boldsymbol{PKP}$ while $s_1, \ldots, s_{N-L}$ are its eigenvalues. We can identify the columns of $\boldsymbol{B}$ with those of $\boldsymbol{A}$ in Equation 8, we have

$$\boldsymbol{B} = \boldsymbol{A} = [\boldsymbol{A}_1, \boldsymbol{A}_2] \tag{16}$$

The eigenvalues of $\boldsymbol{B}_2^{\mathrm{T}}\boldsymbol{K}\boldsymbol{B}_2$ are equivalent to those of $\boldsymbol{PKP}$. The number of non-zero eigenvalues of $\boldsymbol{B}_2^{\mathrm{T}}\boldsymbol{K}\boldsymbol{B}_2$ is $S \leq min(R, N-L)$ so that $s_i = 0$ for $i > S$ and $s_i = \lambda_i$ for $i \leq S$. We also have the relation: $\tilde{\boldsymbol{y}} = \boldsymbol{B}^T\boldsymbol{y}$.

We can now rewrite the restricted log-likelihood from Equation 14 in this setting as:

$$\begin{aligned}
\ell_R(\sigma^2_{quad}, \sigma^2_\epsilon) &= -\frac{1}{2}\left[\sum_{i=1}^{N-L}\log\left(\rho_i\sigma^2_{quad}+\sigma^2_\epsilon\right) + \sum_{i=1}^{N-L}\frac{\tilde{y}_i^2}{\rho_i\sigma^2_{quad}+\sigma^2_\epsilon}\right] + Const \\
&= -\frac{1}{2}\left[\sum_{i=1}^{S}\log\left(\rho_i\sigma^2_{quad}+\sigma^2_\epsilon\right) + (N-L-S)\log\left(\sigma^2_{quad}+\sigma^2_\epsilon\right)\right. \\
&\quad \left. + \sum_{i=1}^{S}\frac{\tilde{y}_i^2}{\rho_i\sigma^2_{quad}+\sigma^2_\epsilon} + \sum_{i=S+1}^{N-L}\frac{\tilde{y}_i^2}{\sigma^2_{quad}+\sigma^2_\epsilon}\right] + Const \\
&= -\frac{1}{2}\left[\sum_{i=1}^{S}\log\left(\rho_i\sigma^2_{quad}+\sigma^2_\epsilon\right) + (N-L-S)\log\left(\sigma^2_{quad}+\sigma^2_\epsilon\right)\right. \\
&\quad \left. - \sum_{i=1}^{S}\frac{\rho_i\sigma^2_{quad}\tilde{y}_i^2}{(\rho_i\sigma^2_{quad}+\sigma^2_\epsilon)(\sigma^2_{quad}+\sigma^2_\epsilon)} + \sum_{i=1}^{N-L}\frac{\tilde{y}_i^2}{\sigma^2_{quad}+\sigma^2_\epsilon}\right] + Const \\
&= -\frac{1}{2}\left[\sum_{i=1}^{S}\log\left(\rho_i\sigma^2_{quad}+\sigma^2_\epsilon\right) + (N-L-S)\log\left(\sigma^2_{quad}+\sigma^2_\epsilon\right)\right. \\
&\quad \left. - \sum_{i=1}^{S}\frac{\rho_i\sigma^2_{quad}\tilde{y}_i^2}{(\rho_i\sigma^2_{quad}+\sigma^2_\epsilon)(\sigma^2_{quad}+\sigma^2_\epsilon)} + \frac{\|\tilde{\boldsymbol{y}}\|_2^2}{\sigma^2_{quad}+\sigma^2_\epsilon}\right] + Const
\end{aligned} \tag{17}$$

Analogously, we can now rewrite the profile restricted log-likelihood from Equation 15 in this setting as:

$$\begin{aligned}
\ell_{PR}(\delta) &= -\frac{1}{2}\left[(N-L)\log\left(\frac{1}{N-L}\left(\sum_{i=1}^{N-L}\frac{\tilde{y}_i^2}{\rho_i+\delta}\right)\right) + \sum_{i=1}^{N-L}\log(\rho_i+\delta)\right] + Const \\
&= -\frac{1}{2}\left[(N-L)\log\left(\frac{1}{N-L}\left(\sum_{i=1}^{S}\frac{\tilde{y}_i^2}{\rho_i+\delta} + \sum_{i=S+1}^{N-L}\frac{\tilde{y}_i^2}{\delta}\right)\right) + \sum_{i=1}^{S}\log(\rho_i+\delta) + (N-L-S)\log(\delta)\right] + Const \\
&= -\frac{1}{2}\left[(N-L)\log\left(\frac{1}{N-L}\left(\sum_{i=1}^{S}\frac{\tilde{y}_i^2}{\rho_i+\delta} + \sum_{i=1}^{N-L}\frac{\tilde{y}_i^2}{\delta} - \sum_{i=1}^{S}\frac{\tilde{y}_i^2}{\delta}\right)\right) + \sum_{i=1}^{S}\log(\rho_i+\delta) + (N-L-S)\log(\delta)\right] + Const \\
&= -\frac{1}{2}\left[(N-L)\log\left(\frac{1}{N-L}\left(-\sum_{i=1}^{S}\frac{\rho_i\tilde{y}_i^2}{(\rho_i+\delta)\delta} + \frac{1}{\delta}\sum_{i=1}^{N-L}\tilde{y}_i^2\right)\right) + \sum_{i=1}^{S}\log(\rho_i+\delta) + (N-L-S)\log(\delta)\right] + Const \\
&= -\frac{1}{2}\left[(N-L)\log\left(\frac{1}{N-L}\left(-\sum_{i=1}^{S}\frac{\rho_i\tilde{y}_i^2}{(\rho_i+\delta)\delta} + \frac{1}{\delta}\|\tilde{\boldsymbol{y}}\|_2^2\right)\right) + \sum_{i=1}^{S}\log(\rho_i+\delta) + (N-L-S)\log(\delta)\right] + Const
\end{aligned} \tag{18}$$

To evaluate the functions in Equations 17 and 18, we need to compute $\tilde{y}_i$ for $i \in \{1, \ldots, S\}$ and $\|\tilde{\boldsymbol{y}}\|_2^2 = \sum_{i=1}^{N-L}\tilde{y}_i^2$.

From Equation 16, we can write $\tilde{\boldsymbol{y}} = \boldsymbol{B}^{\mathrm{T}}\boldsymbol{y} = \begin{bmatrix}\boldsymbol{A}_1^{\mathrm{T}}\boldsymbol{y} \\ \boldsymbol{A}_2^{\mathrm{T}}\boldsymbol{y}\end{bmatrix}$. We see that $\tilde{y}_i$ is given by the $i$-th entry of $\boldsymbol{A}_1^{\mathrm{T}}\boldsymbol{y}$ for $i \leq S$. Here $\boldsymbol{A}_1$ denotes the left singular vectors of $\boldsymbol{P\Phi}$ and can be computed in $\mathcal{O}(ND^2)$ time (as outlined in Section S1.1.2).

Let $\boldsymbol{B}_4 = [\boldsymbol{B}_1, \boldsymbol{B}]$. Since $\boldsymbol{B}_1^{\mathrm{T}}\boldsymbol{B} = \boldsymbol{B}_1^{\mathrm{T}}\boldsymbol{B}_2\boldsymbol{U} = \boldsymbol{0}$, we have:

$$\begin{aligned}
\boldsymbol{B}_4^{\mathrm{T}}\boldsymbol{B}_4 &= [\boldsymbol{B}_1\boldsymbol{B}]^{\mathrm{T}}\begin{bmatrix}\boldsymbol{B}_1 \\ \boldsymbol{B}\end{bmatrix} \\
&= \begin{bmatrix}\boldsymbol{B}_1^{\mathrm{T}}\boldsymbol{B}_1 & \boldsymbol{B}^{\mathrm{T}}\boldsymbol{B}_1 \\ \boldsymbol{B}^{\mathrm{T}}\boldsymbol{B}_1 & \boldsymbol{B}^{\mathrm{T}}\boldsymbol{B}\end{bmatrix} \\
&= \begin{bmatrix}\boldsymbol{I}_L & \boldsymbol{0} \\ \boldsymbol{0} & \boldsymbol{I}_{N-L}\end{bmatrix} = \boldsymbol{I}_N
\end{aligned}$$

Thus, $\boldsymbol{B}_4$ is an orthonormal matrix and we have $\boldsymbol{B}_4\boldsymbol{B}_4^{\mathrm{T}} = \boldsymbol{I}_N$. Further, $\boldsymbol{B}_4^{\mathrm{T}}\boldsymbol{y} = [\boldsymbol{B}_1, \boldsymbol{B}]^{\mathrm{T}}\boldsymbol{y} = \begin{bmatrix}\boldsymbol{B}_1^{\mathrm{T}}\boldsymbol{y} \\ \boldsymbol{B}^{\mathrm{T}}\boldsymbol{y}\end{bmatrix} = \begin{bmatrix}\boldsymbol{y}_1 \\ \tilde{\boldsymbol{y}}\end{bmatrix}$. As a result:

$$\begin{aligned}\|\boldsymbol{y}\|_2^2 = \boldsymbol{y}^{\mathrm{T}}\boldsymbol{y} &= \boldsymbol{y}^{\mathrm{T}}\boldsymbol{I}_N\boldsymbol{y}\\ &= \boldsymbol{y}^{\mathrm{T}}\boldsymbol{B}_4\boldsymbol{B}_4^{\mathrm{T}}\boldsymbol{y}\\ &= \left(\boldsymbol{B}_4^{\mathrm{T}}\boldsymbol{y}\right)^{\mathrm{T}}\boldsymbol{B}_4^{\mathrm{T}}\boldsymbol{y}\\ &= \boldsymbol{y}_1^{\mathrm{T}}\boldsymbol{y}_1 + \tilde{\boldsymbol{y}}^{\mathrm{T}}\tilde{\boldsymbol{y}} = \|\boldsymbol{y}_1\|_2^2 + \|\tilde{\boldsymbol{y}}\|_2^2\end{aligned}$$

We can then compute $\|\tilde{\boldsymbol{y}}\|_2^2 = \|\boldsymbol{y}\|_2^2 - \|\boldsymbol{y}_1\|_2^2$. Computing $\boldsymbol{y}_1$ requires the top $L$ left singular vectors of $\boldsymbol{X}$ that can, in turn, be computed in $\mathcal{O}(NL^2)$ time. To compute $\tilde{y}_i, i \in \{1, \ldots, S\}$, we need to compute eigenvectors (columns of $\boldsymbol{B}$) corresponding to the non-zero eigenvalues of $\boldsymbol{PKP}$ which is equivalent to the corresponding left singular vectors of $\boldsymbol{P\Phi}$. Finally, $\rho_i, i \in \{1, \ldots, S\}$ are the non-zero eigenvalues of $\boldsymbol{PKP}$ which are also obtained from the corresponding singular values of $\boldsymbol{P\Phi}$. Both these quantities can be obtained in $\mathcal{O}(NKD + NK^2 + K^3 + ND^2)$ time. Thus, the profile restricted log-likelihood as represented in Equation 18 can be optimized with a $\mathcal{O}(NKD + NK^2 + K^3 + ND^2)$ one-time computation followed by $\mathcal{O}(D)$ time to evaluate this function subsequently.
